# Structural and Functional Characterization of Host FHL1 Protein Interaction with Hypervariable Domain of Chikungunya Virus nsP3 Protein

**DOI:** 10.1101/2020.09.29.319343

**Authors:** Tetyana Lukash, Tatiana Agback, Francisco Dominguez, Nikita Shiliaev, Chetan Meshram, Elena I. Frolova, Peter Agback, Ilya Frolov

**Author notes:** Corresponding authors: Correspondence and requests for materials should be addressed to I.F. (mailing address: Department of Microbiology, University of Alabama at Birmingham, 1720 2nd Avenue South, BBRB 373/Box 3, Birmingham, AL 35294-2170) or to P.A. (mailing address: Department of Molecular Sciences, Swedish University of Agricultural Sciences, Box 7015, SE-750 07 Uppsala, Sweden).

## Abstract

Decades of insufficient control resulted in unprecedented spread of chikungunya virus (CHIKV) around the globe and millions already suffered from the highly debilitating disease. Nevertheless, the current understanding of CHIKV-host interactions and adaptability of the virus to replication in mosquitoes and mammalian hosts is still elusive. Our new study shows that four-and-a-half LIM domain protein (FHL1) is one of the host factors that interact with hypervariable domain (HVD) of CHIKV nsP3. Unlike G3BPs, FHL1 is not a pre-requisite of CHIKV replication, and many commonly used cell lines do not express FHL1. However, its expression has detectable stimulatory effect(s) on CHIKV replication, and the *Fhl1* KO cell lines demonstrate slower infection spread. The NMR-based studies revealed that the binding site of FHL1 in CHIKV nsP3 HVD overlaps with that of another pro-viral host factor, CD2AP. The structural data also demonstrated that FHL1-HVD interaction is mostly determined by LIM1 domain of FHL1. However, it does not mirror binding of the entire protein, suggesting that other LIM domains are involved. In agreement with previously published data, our biological experiments showed that interactions of CHIKV HVD with CD2AP and FHL1 have additive positive effects on the efficiency of CHIKV replication. This study shows that CHIKV mutants with extensive modifications of FHL1- or both FHL1- and CD2AP- binding sites remain viable and develop spreading infection in multiple cell types. Thus, such modifications of HVD may improve live CHIKV vaccine candidates in terms of their safety and stability of the attenuated phenotype.

**IMPORTANCE:** Replication of chikungunya virus (CHIKV) is determined by a wide range of host factors. Previously, we have demonstrated that the hypervariable domain (HVD) of CHIKV nsP3 protein contains linear motifs that recruit defined families of host proteins into formation of functional viral replication complexes. Now, using NMR-based structural and biological approaches, we have characterized the binding site of cellular FHL1 protein in CHIKV HVD and defined the biological significance of this interaction. In contrast to previously described binding of G3BP to CHIKV HVD, the FHL1-HVD interaction was found to not be a prerequisite of viral replication. However, the presence of FHL1 has a stimulatory effect on CHIKV infectivity and subsequently, the infection spread. FHL1 and CD2AP proteins were found to have overlapping binding sites in CHIKV HVD and additive pro-viral functions. Elimination of FHL1-binding site in nsP3 HVD can be used for the development of stable, live attenuated vaccine candidates.

## INTRODUCTION

Chikungunya virus (CHIKV) is one of the important human pathogens in the *Alphavirus* genus of the *Togaviridae* family (1). In natural circulation, CHIKV is transmitted by mosquito vectors between amplifying vertebrate hosts (2, 3). In mosquitoes, CHIKV develops persistent infection characterized by the high concentration of the virus in salivary glands. Upon infection by mosquito bites, vertebrate hosts develop acute febrile illness characterized by a high level viremia, which is required for transmission to new mosquitoes during the blood meal. Humans can be infected by CHIKV by spillover from the enzootic transmission cycle. However, urban transmissions also became common, and humans may serve as main amplifying hosts with *Aedes aegypti* and *Aedes albopictus* being the transmission vectors (4). In humans, the CHIKV-induced disease is characterized by painful polyarthralgia, fever and rash. In contrast to the diseases caused by other arthritogenic alphaviruses, CHIKV-induced arthralgia can prolong for months to years post infection (PI) (5, 6). Thus, despite lethal outcomes are very rare, CHIKV represents an unquestionable public health threat. Previously, based on its geographical circulation, CHIKV was referred to as the Old World (OW) alphavirus. However, within the last two decades, it has demonstrated an unprecedented spread with wide occurrences in both Old and New Worlds (7).

As for other alphaviruses, the CHIKV genome (G RNA) is represented by a single-stranded RNA of positive polarity of ∼11.5 kb (8). After the release from infectious virions, G RNA serves as a template for translation of four nonstructural proteins, nsP1-4, which are the viral components of replication complexes (vRCs). Initially, nsPs are synthesized as polyprotein precursors P123 and P1234 (9). The partially processed products, P123 and nsP4, mediate the synthesis of the negative-strand RNA to form dsRNA intermediate (10, 11). At later times post infection, the completely processed nsPs function in the transcription of viral G RNA and subgenomic (SG) RNA (12, 13). The latter RNA functions as a template for translation of viral structural proteins, which ultimately package the newly synthesized viral genomes into infectious virions. vRCs reside in the membrane-bound organelles, termed spherules (14). The mechanistical understanding of their assembly and function remains obscure. Their assembly requires participation of a large variety of host proteins, which are indispensable for viral replication. The sets of host factors in vRCs are specific to each alphavirus and particular cell type used in the experiments (15–22).

Nonstructural proteins nsP1, nsP2 and nsP4 have specific enzymatic functions required for the synthesis of viral RNAs and their posttranscriptional modifications (11). NsP3 is an exception, and to date, no direct functions in RNA synthesis have been ascribed for this protein. However, our previous studies and those of other research groups have demonstrated that nsP3 proteins of CHIKV and other alphaviruses are the key determinants of the recruitment of host factors to the sites of vRC assembly at the plasma membrane and to other cytoplasmic complexes, whose functions remain to be better understood (11, 16). Alphavirus nsP3 proteins contain two conserved structured domains (macro domain and alphavirus unique domain [AUD]) (23, 24) and the C-terminal hypervariable domain (HVD). The evolution of the latter domain has been proposed to facilitate adaptation of alphaviruses to hosts and mosquito vector(s) present in specific geographical areas (11, 16). CHIKV nsP3 HVD is intrinsically disordered (25) and contains linear motifs, which interact with cellular proteins (19, 20, 25, 26). Thus, this HVD functions as a hub for assembly of pre-vRCs that contain all CHIKV nsPs, host factors and viral G RNA. The most studied CHIKV HVD-interacting host proteins are members of the G3BP family. G3BP1 and G3BP2 redundantly interact with short repeating peptides located at the C-terminus of the HVD (27, 28). Knockout (KO) of both *G3bp1* and *G3bp2* genes in either rodent or human cells makes them incapable of supporting CHIKV replication (16, 29). Alternatively, the deletion of both elements of the repeat in CHIKV HVD also makes virus nonviable (19). On the other hand, the CHIKV variant with strongly modified HVD, which can interact only with G3BPs, but not with other host factors, is not viable in both vertebrate and mosquito cells (19). This is suggestive that other HVD-interacting cellular proteins are critically involved in CHIKV vRC formation and function. The SH3 domain-containing proteins (CD2AP, BIN1 and SH3KBP1) represent one set of such HVD-interacting partners (19, 20, 26). Binding motifs of the SH3 domain-containing proteins are evolutionarily conserved and are present in HVDs of all of the studied alphaviruses, including those replicating exclusively in insect cells (19, 30). Alterations of these sites have strong negative effects on viral infectivity and viral titers. To date, the binding sites of the SH3 domain-containing proteins, and CD2AP in particular, on CHIKV HVD have been relatively well characterized (19, 25, 26).

Other cellular CHIKV HVD-interacting proteins include the members of FHL and NAPL1 families (19, 20). Thus far, these proteins were demonstrated to interact only with CHIKV HVD and not with the HVDs of other studied alphaviruses. In humans, the family of four-and-a-half LIM domain (FHL) proteins is represented by 4 members, which are expressed in multiple isoforms (31). The FHL1 and FHL2 are the most abandant among family members. Previously, we have identified FHL1 in the co-IP samples of CHIKV HVD isolated from human HEK 293 cells (19), but not from Huh 7 cells. FHL2 has been identified as a CHIKV HVD-interacting partner in mouse NIH 3T3 cells (17). Thus, as it has been shown for G3BP family members and the SH3 domain-containing proteins (25), the FHL1 and FHL2 may redundantly function in CHIKV vRC formation. In this new study, we have applied NMR spectrometry to identify the FHL1-binding sites in CHIKV HVD. In biological experiments, we have developed the cell lines i) expressing no FHL1 protein (*Fhl1* KO) or ii) ectopically expressing GFP-FHL1 fusion (*Fhl1* KO/KI). By using these and other cell and CHIKV variants with mutated binding sites of FHL1 and CD2AP, we demonstrated that the latter proteins are generally dispensable for viral infection but have significant stimulatory effects on CHIKV replication and on viral infectivity. Thus, the FHL proteins, SH3 domain-containing proteins, and likely NAPL1 family members appear to have additive pro-viral effects, but each of them is not critical for CHIKV replication.

## MATERIALS AND METHODS

### Cell cultures

The BHK-21 cells were kindly provided by Paul Olivo (Washington University, St. Louis, MO). Huh 7.5 cells were kindly provided by Charles M. Rice (Rockefeller University, NY). The NIH 3T3, MRC-5, HFF-1, HEK 293T, Hep G2, Vero E6 and HeLa cells were obtained from the American Tissue Culture Collection (Manassas, VA). Additional cell lines, which include HEK 293FT cells and their *Fhl1* KO derivative were kindly provided by Dr. Ali Amare (32). Huh 7.5, MRC-5 and HFF-1 cells were maintained in Dulbecco’s modified Eagle medium supplemented with 10% fetal bovine serum (FBS). Other cell lines were maintained in alpha minimum essential medium supplemented with 10% FBS and vitamins.

### Plasmid constructs

Plasmid encoding infectious cDNA of CHIKV 181/25 containing GFP under control of viral subgenomic promoter was described elsewhere (33). To introduce mutations into HVD, gene blocks from Integrated DNA Technologies (IDT) were cloned into cDNA of CHIKV 181/25 to replace the original wt nucleotide sequence. The presence of correct mutations was verified by sequencing of HVD in the final constructs. Plasmid encoding CHIKV/nsP3-Cherry was described elsewhere (16). Plasmids encoding VEEV replicons with mutated CHIKV HVDs fused with Flag-GFP under control of the subgenomic promoter had the previously described design (16). For the development of *Fhl1* and *Fhl2* KO cell lines, guide RNAs (gRNAs) were designed to target nucleotide (nt) sequences close to the initiating codons of the indicated genes to prevent the synthesis of truncated proteins with possible unknown functions. gRNA sequences were cloned into modified GeneArt CRISPR Nuclease [CD4 Enrichment] vector, which also encodes Cas9 and CD4 genes (Life Technologies). *hFhl1* gene encoding FHL1A isoform fused with GFP was cloned into modified PiggyBac plasmid (System Bioscience, Inc) under control of the CMV promoter. This plasmid was designed to express bi-cistronic mRNA, which encoded FHL1-GFP and blasticidin resistance gene cloned under control of the internal ribosome entry site of encephalomyocarditis virus (EMCV IRES). The codon-optimized sequences of the full-length FHL1 and individual LIM domains (LIM1 aa 1-97, LIM2 aa 96-157, LIM3 aa157-220, LIM4 aa 213-280) were obtained as synthetic DNA fragments from IDT and cloned into pE-SUMOpro-3 plasmid (LifeSensors Inc.) as fusions with SUMO.

### Generation of KO and KI cell lines

gRNAs-encoding plasmids were transfected into HEK 293T, HeLa and NIH 3T3 cells using TransIT-X2 reagent (Mirus). At 2 days post transfection, cells expressing the surface CD4 receptor were enriched using the Dynabeads CD4 cell isolation kit (Life Technologies). They were seeded to develop single cell-derived clones. Clones were analyzed for the lack of expression of targeted proteins by Western Blot (WB), and for each cell line, 2**-**3 clones demonstrating the complete absence of targeted proteins were further analyzed. The targeted fragments in cellular genomes were amplified by PCR and cloned into pRS2 plasmid. Multiple clones were sequenced to confirm that both alleles were mutated. To generate a stable FHL1-GFP-expressing cell line, the above-described PiggyBac-derived plasmid was transfected into KO *Fhl1* HEK 293T cell line using TransIT-X2 reagent (Mirus). Next day, cells were re-seeded at different densities and incubated in the media supplemented with blasticidin (10 μg/ml) to develop colonies. Cell clones demonstrating the lowest detectable levels of GFP were selected for further experiments.

### Rescuing of the viruses

Plasmid DNAs encoding cDNAs of viral genomes were purified by ultracentrifugation in CsCl gradients. DNAs were linearized using NotI restriction site located immediately downstream of poly(A) tail. RNAs were *in vitro*-synthesized by SP6 RNA polymerase (New England Biolabs) in the presence of a cap analog (New England Biolabs) according to the manufacturer’s recommendations. The qualities of the synthesized RNAs were tested by electrophoresis in nondenaturing agarose gels, and aliquots of transcription reactions containing equal amounts of RNAs were used for electroporation without additional purification. Electroporations of BHK-21 cells by *in vitro*-synthesized RNAs were performed under previously described conditions (34, 35). Viruses were harvested at 24 h post electroporation, and infectious titers were determined by a plaque assay on BHK-21 cells (36). To rule out a possibility that the designed CHIKV/GFP mutants require adaptive mutations for their viability, the infectious center assay (ICA) was applied as an additional control. The defined numbers of electroporated cells were seeded on the subconfluent monolayers of naïve BHK-21 cells in 6-well Costar plates. After cell attachment, monolayers were covered by 0.5% agarose, supplemented in DMEM supplemented with 3% FBS. After ∼60 h of incubation at 37°C, plaques were stained with crystal violet.

### Western blot analyses

Equal numbers of cells indicated in the figures were lysed directly in protein gel loading buffer. Proteins were separated in either 4–12% NuPAGE gels (ThermoFisher) or in custom 10% polyacrylamide gels and transferred to nitrocellulose membrane. The following antibodies were used for detection of cellular proteins: anti-FHL1 (10991-1-AP, Proteintech), anti-FHL2 (21619-1-AP, Proteintech), anti-GAPDH antibodies (6ooo4-1-Ig, Proteintech) and anti-β-actin (66009-1-Ig, Proteintech). Secondary Abs labeled with IRDye 680RD or IRDye 800CW infrared dyes were acquired from LI-COR Biosciences. Membranes were scanned and analyzed on LI-COR imager.

### Viral infections

For comparative analyses of viral replication, equal numbers of different cells indicated in the figures were seeded into the wells of 6-well Costar plates. After 4 h of attachment at 37°C, cells were infected at MOIs indicated in the figure legends in phosphate buffered saline (PBS) supplemented with 1% FBS. After incubation for 1 h at 37°C, cells were washed with the media and then incubated for the times indicated in figure legends. Viral titers in the harvested samples were determined by plaque assay on BHK-21 cells.

### RT-qPCR

Total RNAs were extracted from the cells indicated in the figures using Direct-zol RNA MiniPrep kit according to the manufacturer’s instructions (Zymo Research). Equal amounts of RNA samples were used for cDNA synthesis by QuantiTect reverse transcription (RT) kit according to the manufacturer’s instructions (Qiagen). qPCR was performed using the following primers: i) forward 5’TACTGCGTGGATTGCTAC3’ and reverse 5’CCAGGATTGTCCTTCATAG3’ (human *Fhl1*), ii) forward 5’ACTGCTTCTGTGACTTGTATG3’ and reverse 5’AAGCAGTCGTTATGCCAC3’ (human *Fhl2*), iii) forward 5’ATTACTGCGTGGATTGCTAC3’ and reverse 5’CTTCATAGGCCACCACAC3’ (mouse *Fhl1*), iv) forward 5’TCCTCACAGAGAGAGATGAC3’ and reverse 5’CTGGTTATGAAAGAAAACATGCG3’ (mouse *Fhl2*). qPCR reactions were performed using SsoFast EvaGreen supermix (Bio-Rad) in a CFX96 real-time PCR detection system (Bio-Rad). The specificities of the qPCR products were confirmed by analyzing their melting temperatures. The data were normalized to the mean threshold cycle (CT) of 18S ribosomal RNA in each sample. The fold differences were calculated using the ΔΔCT method.

### Viral infectivities

Equal numbers of cells of different cell lines were seeded into the wells of 6-well Costar plates. After 4 h of incubation at 37°C, different cell lines were infected at the same time with the same serial 10-fold dilutions of the viral stock for 1 h at 37°C. Cells were covered with 0.5% agarose or Avicel RC-591NT supplemented with DMEM and 3% FBS. Foci of GFP-positive cells were imaged at either 48 or 24 h PI. The cell line-specific titers were determined in GFP-positive foci-forming infectious units (inf.u/ml).

### Immunostaining

Cells were seeded into 15 μ-Slide 8-well plates (Ibidi) (2×10^4^ cells/well) and infected at MOIs indicated in the figure legends. At the indicated times, cells were fixed with 4% paraformaldehyde (PFA), permeabilized and stained with following Abs: anti-CHIKV nsP3 (MAB2.19, custom), anti-FHL1 (10991-1-AP, Proteintech), anti-FHL2 (21619-1-AP, Proteintech), anti-CD2AP (sc-9137, Santa Cruz Biotechnology), anti-SH3KBP1 (sc-166862, Santa Cruz Biotechnology) and anti-G3BP1 (kindly provided by Richard Lloyd). After staining with fluorescent secondary Abs, images were acquired on a Zeiss LSM 800 confocal microscope in Airyscan mode with a 63X 1.4NA PlanApochromat oil objective.

### Protein purification

Unlabeled and ^15^N-labeled CHIKV HVD (aa 325–523) of strain 181/25 was expressed as a fusion with Twin-Strep-tag (GSWSHPQFEKGGGSGGGSGGGSWSHPQFEK) from pE-SUMOpro-3 plasmid (LifeSensors Inc.) in E. coli strain Rosetta 2 (DE3) (Novagen) as described elsewhere (25). LIM1, LIM2, LIM3, LIM4 domains and the full-length hFHL1 were produced in M9 media supplemented with 2 g/L of NH_4_Cl and 3 g/L of glucose in E. coli LEMO21(DE3) (New England Biolabs) or Rosetta2(DE3)pLacI (Novagen). Protein expression was induced by 1mM IPTG after cells reached the density of ∼1 OD600, and then cells were incubated at 20°C for 20 h. Freshly prepared or frozen pellets were lysed in Emulsiflex B15 (Avestin). The recombinant proteins were purified on a HisTrap HP column (GE Healthcare), followed by buffer exchange on a HiPrep^TM^ 26/10 desalting column (GE Healthcare). After cleavage of SUMO tag, proteins were additionally purified by passing through HisTrap HP column (GE Healthcare). Size exclusion chromatography on a Superdex 75 10/300 column (GE Healthcare) in phosphate buffer [50 mM N_2_HPO_4_ pH 6.8, 200 mM NaCl, 2 mM tris(hydroxypropyl)phosphine (TCEP)] was used as a final purification step. Purified proteins were diluted to a final concentration of 50 mM NaCl and then concentrated on Amicon centrifugal filters 30K for FHL and 10K for LIM domains (Millipore). The proteins’ purities and identities were confirmed by SDS-PAGE and mass spectrometry. Protein concentrations were determined on 280 nm using an extinction coefficient of 22,760 M^−1^cm^−1^ (CHIKV HVD), 38,120 M^−1^cm^−1^ (FHL1), 9,530 M^−1^cm^−1^ (LIM1), 8,250 M^−1^cm^−1^ (LIM2), 10,810 M^−1^cm^−1^ (LIM3) and 10,810 M^−1^cm^−1^ (LIM4), which were determined using ProteinCalculator v3.4 (http://protcalc.sourceforge.net/).

### NMR titration experiments

Binding experiments were performed by recording 2D best-TROSY (^1^H -^15^N) spectra on Bruker Avance III spectrometer equipped with a cryo-enhanced QCI-P probe and operating at 600.18MHz for ^1^H and 60.82MHz for ^15^N nuclei. The spectra were recorded with acquisition times of 109.6 and 141.9 ms in (*t_1_, t_2_*) in 200 x 1024 complex matrices. Hard ^1^H pulses were applied with 27.5 a kHz field and ^15^N pulses with a 7.14 kHz field, carriers were positioned at 4.68 and 118ppm for ^1^H and ^15^N, respectively. A relaxation delay of 0.3 s was employed, and 32 scans were collected. The control 1D spectra were used to assess the purity and stability of the samples. 0.1 mM of sodium trimethylsilylpropanesulfonate (DSS) was added as a reference for chemical shift. All NMR spectra were processed by TopSpin4.0.6 software.

For processing spectra, a Gaussian window function with parameters LB=-10 and GB=0.1 in F2 and GB=0.3 in F1 dimensions was used to obtain the 2K x 1K spectra with linear prediction in F1. The unlabeled LIM1, LIM2, LIM3, LIM4 and FHL1 proteins were titrated into uniformly ^15^N-labelled CHIKV HVD samples at 25°C.

Analyses of the spectra were performed using CcpNmr software package 2.4.2. Intensities/or volumes of cross-peaks were extracted by Gaussian /or peak fit methods, respectively. The presented data were normalized to the apo form intensities. The magnitudes of the combined chemical shift deviations (Δδ_(H+N)_) were calculated using equation 1 (37):

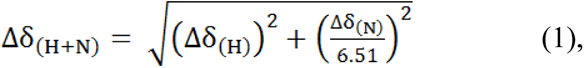

in which the proton (Δδ_(H)_) and the nitrogen (Δδ_(N)_) chemical shift perturbations (CSP) were defined as a difference in chemical shift between the equilibrium mixture of the bound (δ^1^H_bound_ and δ^15^N_bound_, correspondingly) and the free states (δ^1^H_free_ and δ^15^N_free_, correspondingly) induced by different proteins (eq. 2).

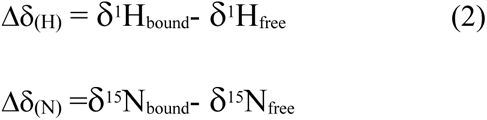

## RESULTS

### Mammalian cell lines express different levels of FHL1 and FHL2

In the previous studies, we analyzed the composition of protein complexes formed on CHIKV HVD during its expression in vertebrate cells as a Flag-GFP-HVD fusion (16, 17, 19). In the NIH 3T3 and HEK 293T cells, these HVD complexes contained members of G3BP and NAPL1 families, SH3 domain-containing proteins and readily detectable levels of FHL2 or FHL1, respectively. We did not detect any members of FHL family in CHIKV HVD co-IP samples derived from Huh 7 cells (19). These results suggested that the members of the FHL family might function in CHIKV replication. Therefore, since in different cells CHIKV HVD complexes contained only FHL1, FHL2 or neither, we analyzed expression of FHL1 and FHL2 proteins in several mammalian cell lines. Expression levels of FHL3 were not investigated because of its absence in CHIKV HVD protein complexes in the previous mass spectrometry-based studies. The cell lines analyzed included HEK 293T, Huh 7.5, Hep G2, MRC-5, HFF-1, NIH 3T3, BHK-21 and Vero E6 cells (Fig. 1A). Both FHL1 and FHL2 proteins were most efficiently detected in HEK 293T and HFF-1 cells. MRC-5 and BHK-21 cells also demonstrated readily detectable levels of FHL1 and relatively high levels of FHL2. In Vero E6, NIH 3T3, Huh 7.5 and Hep G2 cells, FHL2 was definitively present, albeit its concentrations in Huh 7.5 and Hep G2 were low. Interestingly, in the latter cell lines, FHL1 was undetectable by Western blotting (WB). Since WB is not a sensitive method for gene expression analysis, next we used qRT-PCR to evaluate concentration of specific mRNAs in some of the cells. The qPCR data (Fig. 1B) were in agreement with the WB results. Huh 7.5 and Hep G2 cells did not contain detectable levels of FHL1-specific mRNA, and the levels of FHL2 mRNA were relatively low compared to those found in HEK 293T cells. MRC-5 cells contained high levels of both FHL1- and FHL2-specific mRNAs. Murine NIH 3T3 cells also had no FHL1 mRNA, but FHL2 mRNA was found at a readily detectable level. In agreement with published data (31), FHL1 mRNA concentrations were higher in the control samples of mouse heart and muscle tissues, while FHL2 mRNA was absent in these muscles.

**FIG. 1.**
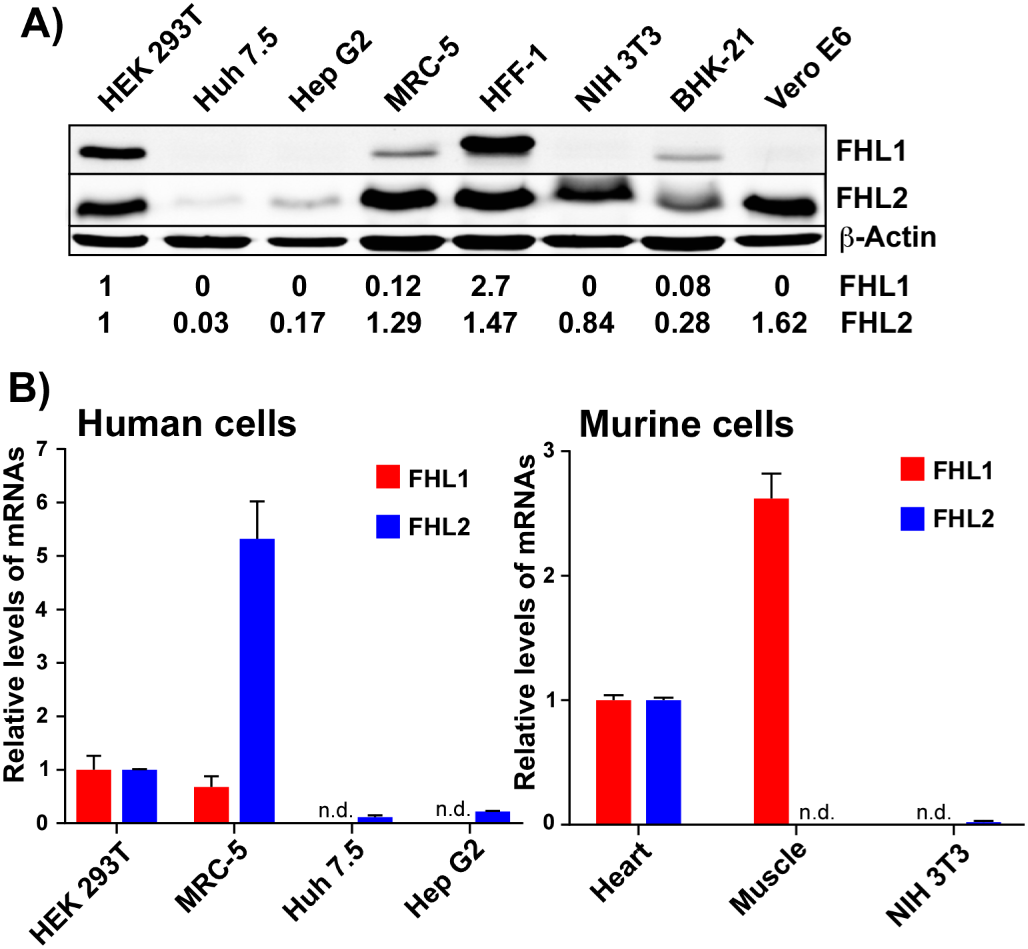
Vertebrate cell lines strongly differ in the levels of FHL1 and FHL2 expression. A) Lysates of different cell lines were analyzed by WB for the expression of FHL1 and FHL2 proteins using specific Abs. The membranes were scanned on LI-COR imager. Relative levels of proteins were determined using the manufacturer’s software. The data were normalized to the levels of β-actin and the levels of FHL1 and FHL2 proteins in HEK 293T cells. B) Concentration of FHL1- and FHL2-specific mRNAs in the indicated cells. The data were normalized to the levels of 18S ribosomal RNA in each sample. For human and mouse cells, the data were also normalized to concentrations of the mRNAs in HEK 293T and mouse heart muscle, respectively. n.d. indicates nondetectable.

Taken together, these data demonstrated that the commonly used cell lines strongly differ in the expression of FHL1 protein, but FHL2 is more constitutively expressed. Huh 7.5, Hep G2 and NIH 3T3 cells do not express *Fhl1* at protein and mRNA levels and thus, can be used in the experiments as natural analogs of *Fhl1* KO cells.

### The cell lines that express no FHL1 can support CHIKV replication

Since FHL1 was expressed at detectable levels in relatively few cell lines (Fig. 1), we tested whether the FHL1-negative cells could support CHIKV replication. In these and further experiments, we used recombinant CHIKV 181/25 (38) with the GFP gene cloned under control of the duplicated subgenomic promoter (CHIKV/GFP) (33). GFP expression was used to monitor the numbers of infected cells and the infection spread over time. The used CHIKV 181/25 strain has been generated by serial passaging of CHIKV AF15561 in cultured cells (38, 39). This resulted in adaptation of E2 glycoprotein to bind heparan sulfate for the entry into different cells and thus, made infection less dependent on one of the natural CHIKV receptors, which is not expressed in the cell lines, such as HEK 293T and HeLa cells (40). Importantly, the attenuated phenotype of CHIKV 181/25 is determined by two point mutations in E2 glycoprotein (38), while the encoded nsP3 is exactly the same as in the parental CHIKV strain AF15561 (39). The nonstructural polyprotein of CHIKV 181/25 contains only a single conservative aa substitution in nsP1, which has no effect on viral replication and pathogenesis (38). Thus, the data concerning the interaction of CHIKV 181/25-based nsPs with host factors and vRC formation are fully applicable to the parental pathogenic strain.

In agreement with the previously published data (41–44), despite lack of FHL1 expression, both Huh 7.5 and Hep G2 cells supported CHIKV/GFP replication (Figs. 2A and B). In repeat experiments, titers of the samples harvested from Huh 7.5 cells were always detectably higher than those in the samples harvested from Hep G2. They generally correlated with the inability of CHIKV/GFP to infect the Hep G2 monolayers completely, while 100% of Huh 7.5 were infected by 24 h PI (Fig. 2C). Typically, almost half of Hep G2 cells remained negative in terms of GFP expression and thus, viral replication. These cells survived the infection even after prolonged incubation in the presence of high levels of the released virus. The reason for this partial cell resistance remains unclear. However, this is not a result of type I IFN accumulation in the media, because at 24 h PI, concentration of the released IFN-β remained below the levels of detection (data not shown). The above data (Fig. 1) demonstrated that Hep G2 and Huh 7.5 cells have the same phenotype in terms of lack of FHL1 expression and low level of FHL2. However, the Huh 7.5 cells were highly susceptible to CHIKV/GFP infection and developed complete cytopathic effect (CPE) by 48 h PI. They produced CHIKV/GFP essentially as efficiently as BHK-21 cells, but some delay in infectious virus release was reproducibly detected at the early times PI (Fig. 2A and B).

**FIG. 2.**
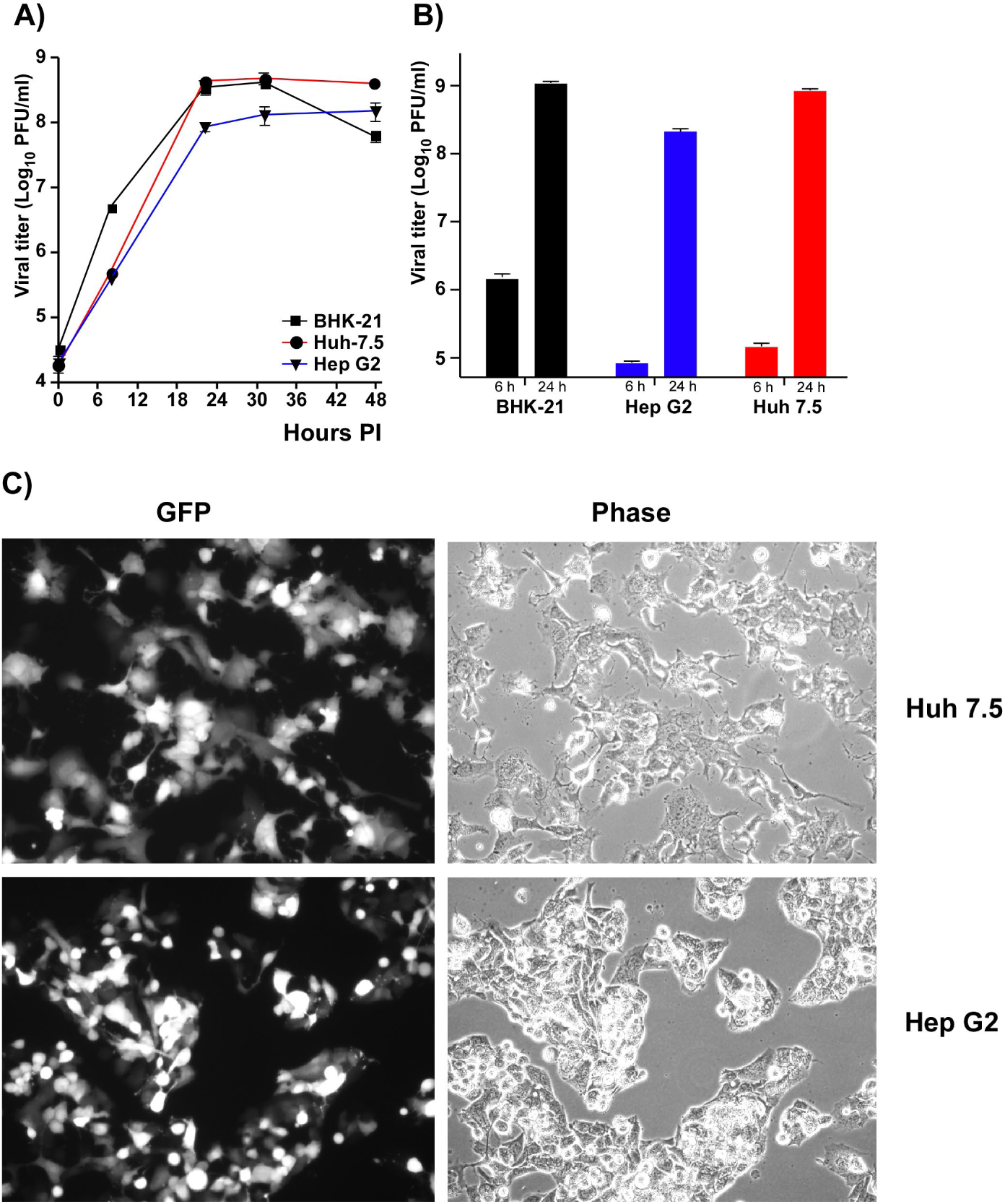
The cell lines expressing no FHL1 can support CHIKV replication. A) 5×10^5^ cells of the indicated cell lines were seeded into 6-well Costar plates. After 3 h of incubation at 37°C, they were infected with CHIKV/GFP at an MOI of 10 PFU/cell as described in Materials and Methods. Media were harvested at the indicated time points, and viral titers were determined by plaque assay on BHK-21 cells. This experiment was repeated 2 times with the same results. B) The indicated cell lines were infected at an MOI of 1 PFU/cell and titers of the released virus were assessed at 6 and 24 h PI by plaque assay on BHK-21 cells. The experiment was repeated 3 times. Means and standard deviations (SDs) are presented. C) Images of infected Huh 7.5 and Hep G2 cells in the experiment presented in panel B, were acquired at 24 h PI on the fluorescence microscope.

Taken together, these data and the results of the previous studies on NIH 3T3 cells (45), which do not express FHL1 (Figs. 1A and B), demonstrated that expression of the latter protein is not a prerequisite of CHIKV replication at least in some of the commonly used human and murine cell lines.

### *Fhl1* KO and *Fhl2* KO cells can support CHIKV replication

The above experiments did not rule out a possibility that members of the FHL family have pro-viral activity. Therefore, to further evaluate the role(s) of FHL1 for CHIKV replication, we developed *Fhl1* KO derivatives of HEK 293T and HeLa cells using the CRISPR/Cas9 approach (see Materials and Methods for details). The cloned cell lines were characterized for the lack of FHL1 expression and alteration of open reading frame in the first exon in the *Fhl1* gene. In the selected clones of both cell lines, no FHL1 expression was detected by WB using FHL1-specific Abs (Fig. 3A), and both *Fhl1* KO HeLa and *Fhl1* KO HEK 293T cells contained insertions of a single nucleotide at the same position of the target sequence in both alleles (Fig. 3B).

**FIG. 3.**
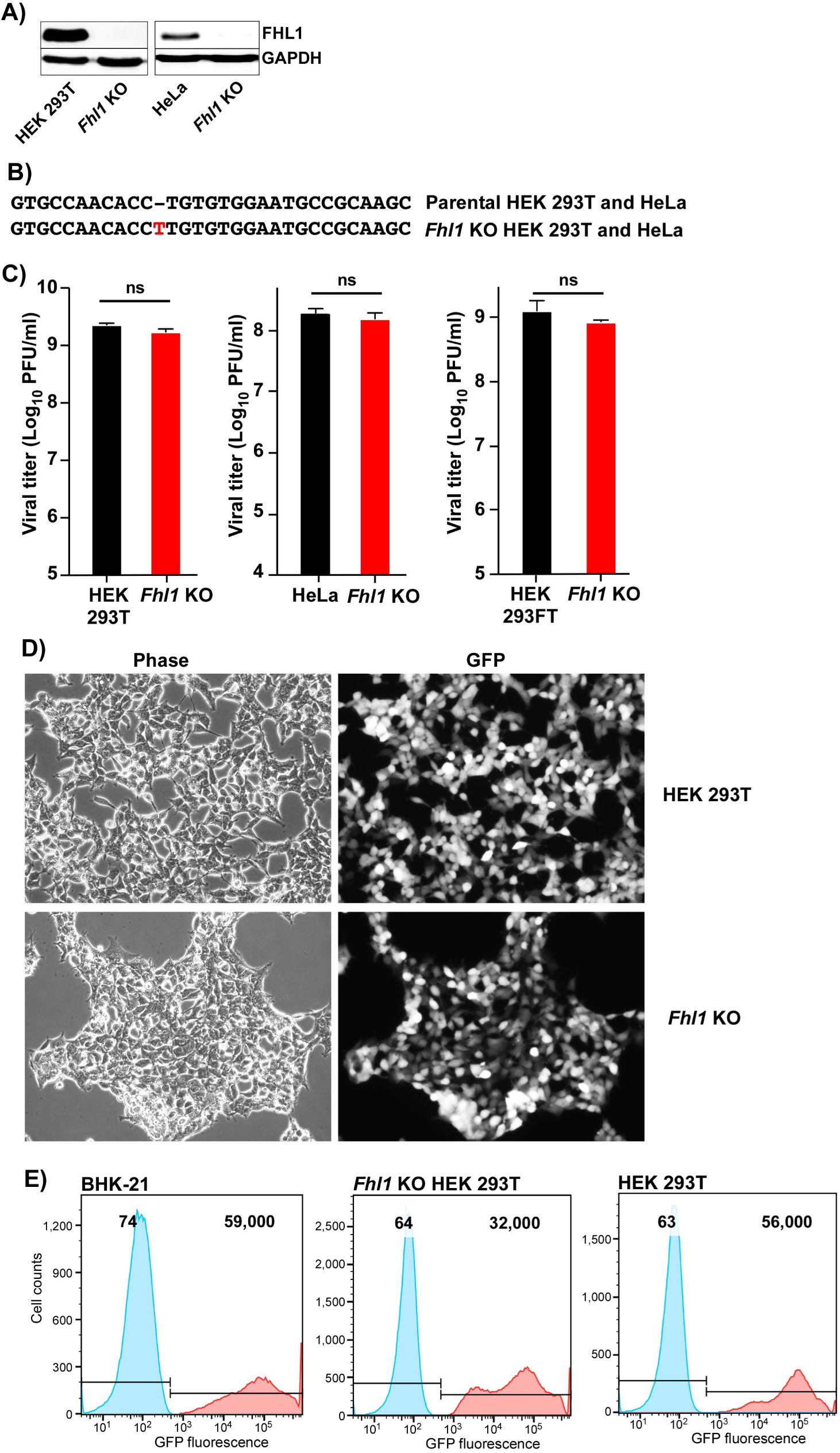
*Fhl1* KO cells support CHIKV replication. A) The results of WB analysis of FHL1 expression in HEK 293T, *Fhl1* KO HEK 293T, HeLa and *Fhl1* KO HeLa cells. B) Mutation identified in both alleles of *Fhl1* KO HEK 293T and *Fhl1* KO HeLa cells. C) The indicated cell lines in 6-well Costar plates were infected with CHIKV/GFP at an MOI of 1 PFU/cell and viral titers were assessed at 24 h PI by plaque assay on BHK-21 cells. This experiment was repeated 3 times and means and SDs are presented. The two-way ANOVA Sidak test was used for statistical analysis. ns indicates nonsignificant. D) Images of infected KO *Fhl1* HEK 293T and parental HEK 293T cells were acquired at 24 h PI on the fluorescence microscope in the experiment presented in panel C. E) The indicated cell lines were infected with CHIKV/GFP at an MOI of 10 PFU/cell and at 24 h PI, analyzed for GFP expression by flow cytometry. Control mock-infected cells are shown in blue, and the corresponding infected cells-in red. Numbers indicate mean fluorescence intensities.

The KO cell lines were infected with CHIKV/GFP at an MOI of 1 PFU/cell, and surprisingly, in contrast to the previously published data (32), in our experimental conditions, the effect of *Fhl1* KO on viral replication was undetectable. In repeat experiments, at this MOI, the KO cells supported CHIKV/GFP replication to the same titers as the parental HEK 293T cell line (Fig. 3C) and demonstrated high levels of GFP expression (Fig. 3D). Similarly, no difference in viral titers was detected between HeLa cells and their *Fhl1* KO derivative (Fig. 3C). Here we present the data generated on a single clone of each cell line, but other tested KO clones produced identical results, suggesting that the lack of the KO effect was not cell clone-specific. At higher MOI (10 PFU/cell), by 24 h PI, *Fhl1* KO HEK 293T cells were not only all GFP positive (Fig. 3E), but also demonstrated signs of CPE (data not shown). These results strongly differed from those previously published on another *Fhl1* KO cell line (32). Therefore, we repeated our experiments on the previously described *Fhl1* KO cells and their parental counterpart (HEK 293FT *Fhl1* KO and HEK 293FT, respectively), which were kindly provided by the authors. Despite HEK 293FT *Fhl1* KO cells contained an open reading frame (ORF) shift in different exon, we observed the same results. At an MOI of 1 PFU/cell, by 24 h PI, the KO cells released CHIKV/GFP to the same high titer as the parental counterpart (Fig. 3C).

Next, we analyzed accumulation of known HVD-binding host proteins in cytoplasmic nsP3 complexes formed in infected *Fhl1* KO cells (Fig. 4). The nsP3 complexes certainly did hot contain FHL1. However, we detected a low-level FHL2 accumulation. This observation correlated with the previously described interaction of mouse FHL2 with CHIKV HVD in NIH 3T3 cells (17). It suggests that FHL2 might at least partially substitute for FHL1 in the absence of the latter protein in human cells. In *Fhl1* KO cells, the SH3 domain-containing proteins, which include CD2AP and SH3KBP1, continued to accumulate in CHIKV nsP3- and G3BP1-containing complexes.

**FIG. 4.**
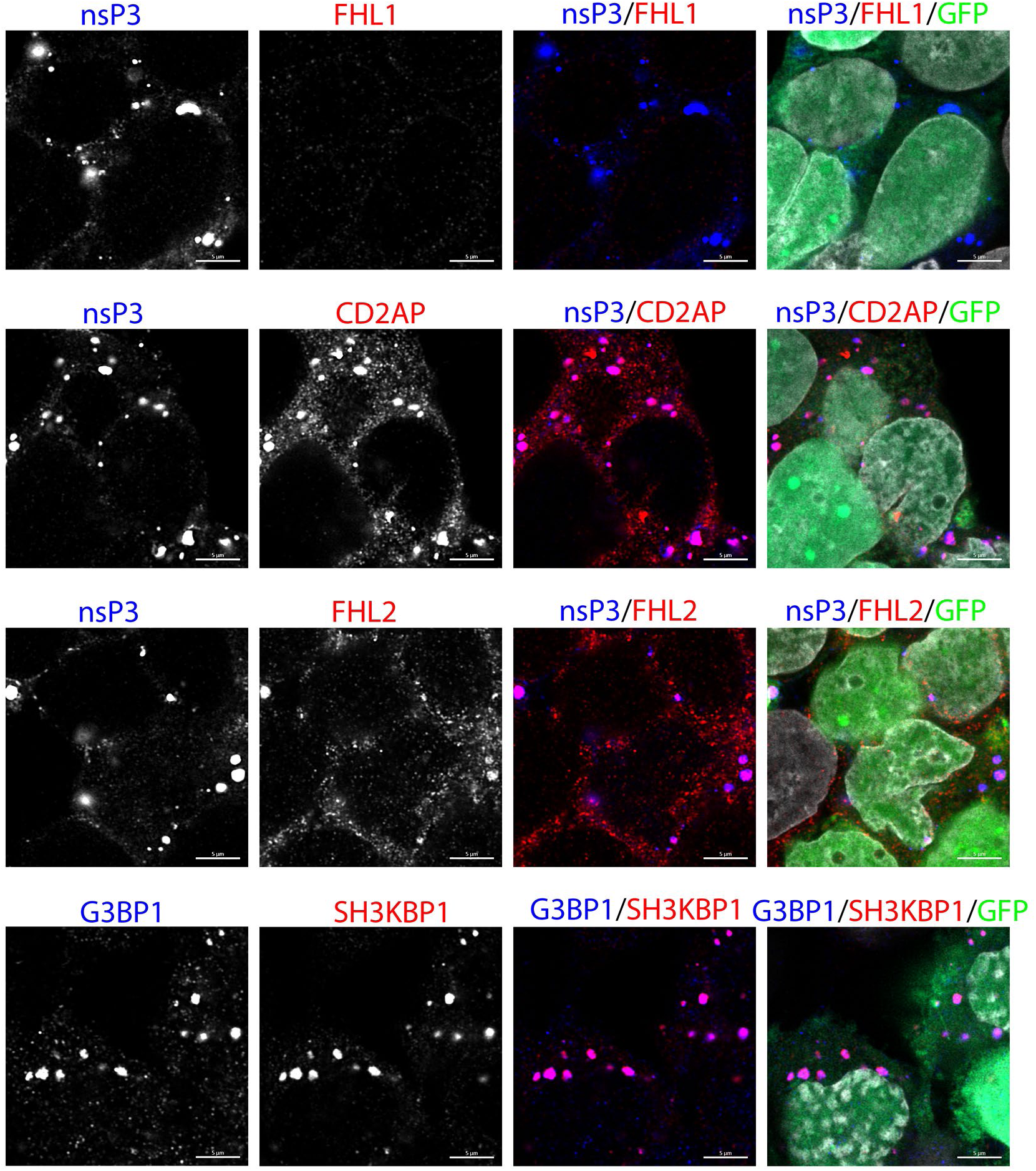
The cytoplasmic CHIKV nsP3 complexes are formed in the *Fhl1* KO HEK 293T cells and contain host proteins. *Fhl* KO HEK 293T were infected with CHIKV/GFP at an MOI of 1 PFU/cell and fixed at 15 h PI with 4% PFA. They were immunostained with antibodies specific to nsP3, FHL1, CD2AP or FHL2. On the bottom panel, cells were stained with antibodies against G3BP1, which is the marker for cytoplasmic nsP3 complexes, and SH3KBP1. Images were acquired on a Ziess LSM800 microscope in Airyscan mode. All but FHL1 proteins were colocalized with nsP3 complexes. Scale bar is 10 μm.

Since the results from our previous studies showed the presence of FHL2 in CHIKV HVD complexes isolated from NIH 3T3 cells (17), next, we intended to evaluate the effect of murine FHL2 expression and generated an *Fhl2* KO derivative of NIH 3T3 cell line. Sequencing of the target fragment in the genome of the selected clone demonstrated the insertion of a single nucleotide into both alleles, which altered ORF starting from the first exon (Fig. 5A). Lack of FHL2 expression was confirmed by WB using FHL2-specific Abs (Fig. 5B). We also analyzed a possibility that KO of *Fhl2* could induce expression of FHL1, as this phenomenon was previously detected during generation of single *G3bp* KOs. No induction of FHL1 was found, and *Fhl2* KO NIH 3T3 cells did not express either FHL1 or FHL2 proteins (Fig. 5B).

**FIG. 5.**
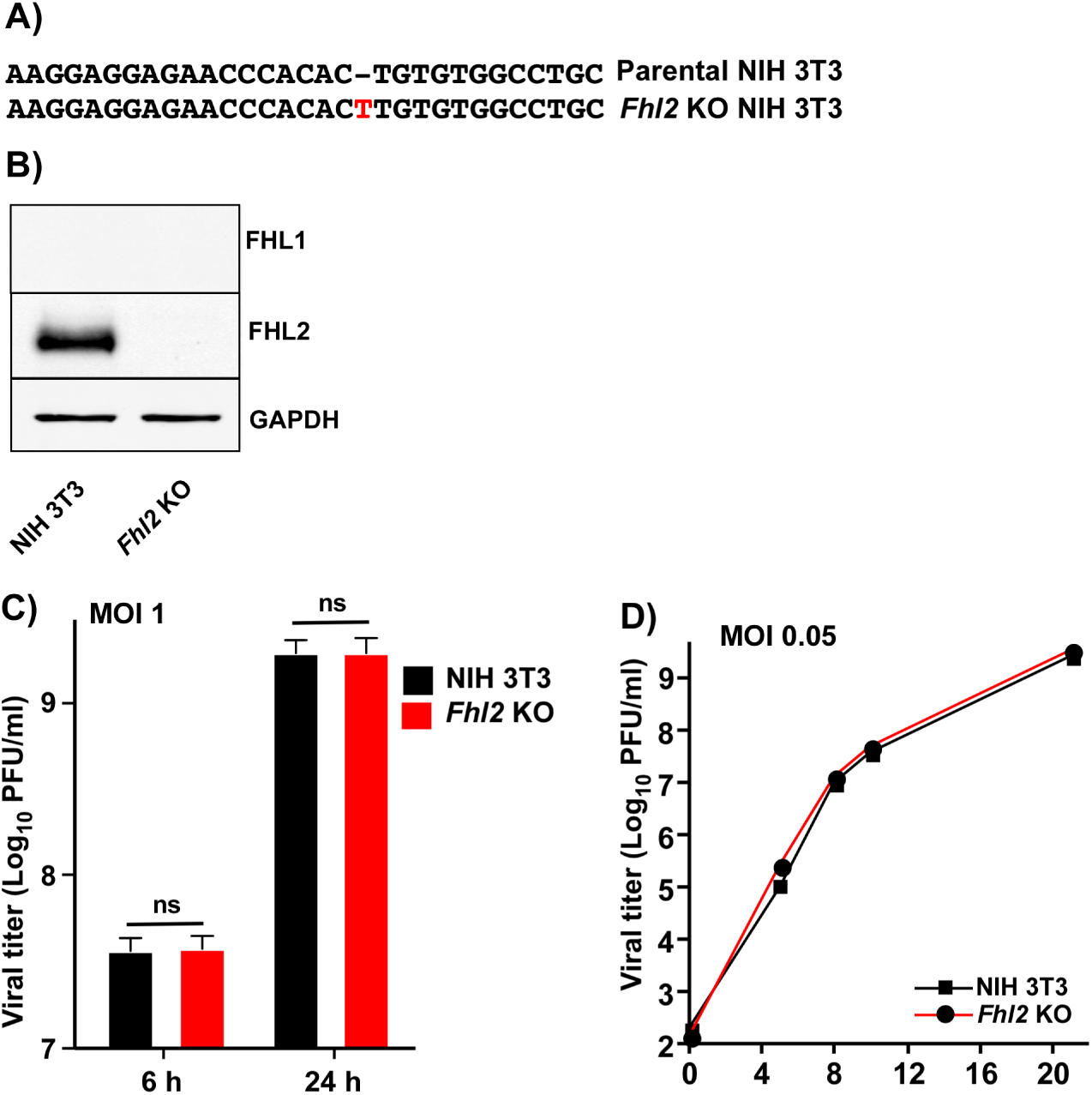
*Fhl2* KO NIH 3T3 cells, which express no FHL1 and FHL2, support efficient CHIKV replication. A) Mutation identified in both alleles of *Fhl2* KO NIH 3T3 cells. B) The results of WB analysis of FHL1 and FHL2 expression in KO *Fhl2* NIH 3T3 cells. C) NIH 3T3 cells and their KO derivatives in 6-well Costar plate were infected with CHIKV/GFP at an MOI of 1 PFU/cell. Viral titers were analyzed at 6 and 24 h PI, by plaque assay on BHK-21 cells. The experiment was repeated 3 times and means and SDs are presented. The two-way ANOVA Sidak test was used for statistical analysis. ns indicates nonsignificant. D) The indicated cell lines were infected with CHIKV/GFP at an MOI of 0.05 PFU/cell. Samples were harvested at the indicated times PI, and titers were assessed by plaque assay on BHK-21 cell. The experiment was reproducibly repeated twice, and the results of one of them are presented.

Next, the *Fhl2* KO and parental NIH 3T3 cells were infected as above with CHIKV/GFP at an MOI of 1 PFU/cell and viral titers were assessed at 6 and 24 h PI (Fig. 5C). At 24 h time point, all of the cells in both cell lines were infected, demonstrated high levels of GFP expression, and exhibited profound CPE (data not shown). Either at 6 or 24 h PI, no differences in viral titers were detected. No difference in viral replication was also found at lower MOI of 0.05 PFU/cell (Fig. 5D). Based on these data, we concluded that *Fhl2* KO NIH 3T3 cells could support CHIKV/GFP replication even in the absence of expression of both FHL1 and FHL2.

Taken together, the data from these experiments and from our previous studies of CHIKV replication (19) suggested that, as we described for the SH3 domain-containing proteins, FHL family members are not as critical for CHIKV replication as G3BP family members and are generally dispensable.

### FHL1 protein has pro-viral activity

Next, we analyzed the ability of CHIKV/GFP to develop a spreading infection in HEK 293T and KO *Fhl1* derivative. The identical monolayers were infected with a few PFUs of CHIKV/GFP followed by incubation in the media supplemented with 0.5% Avicel (see Materials and Methods for details). At 24 h PI, the GFP-positive foci formed on KO *Fhl1* cells were significantly smaller than those formed by the same virus on parental HEK 293 cells (Fig. 6A). This was the first indication that a lack of FHL1 expression has negative effect on CHIKV infection.

**FIG. 6.**
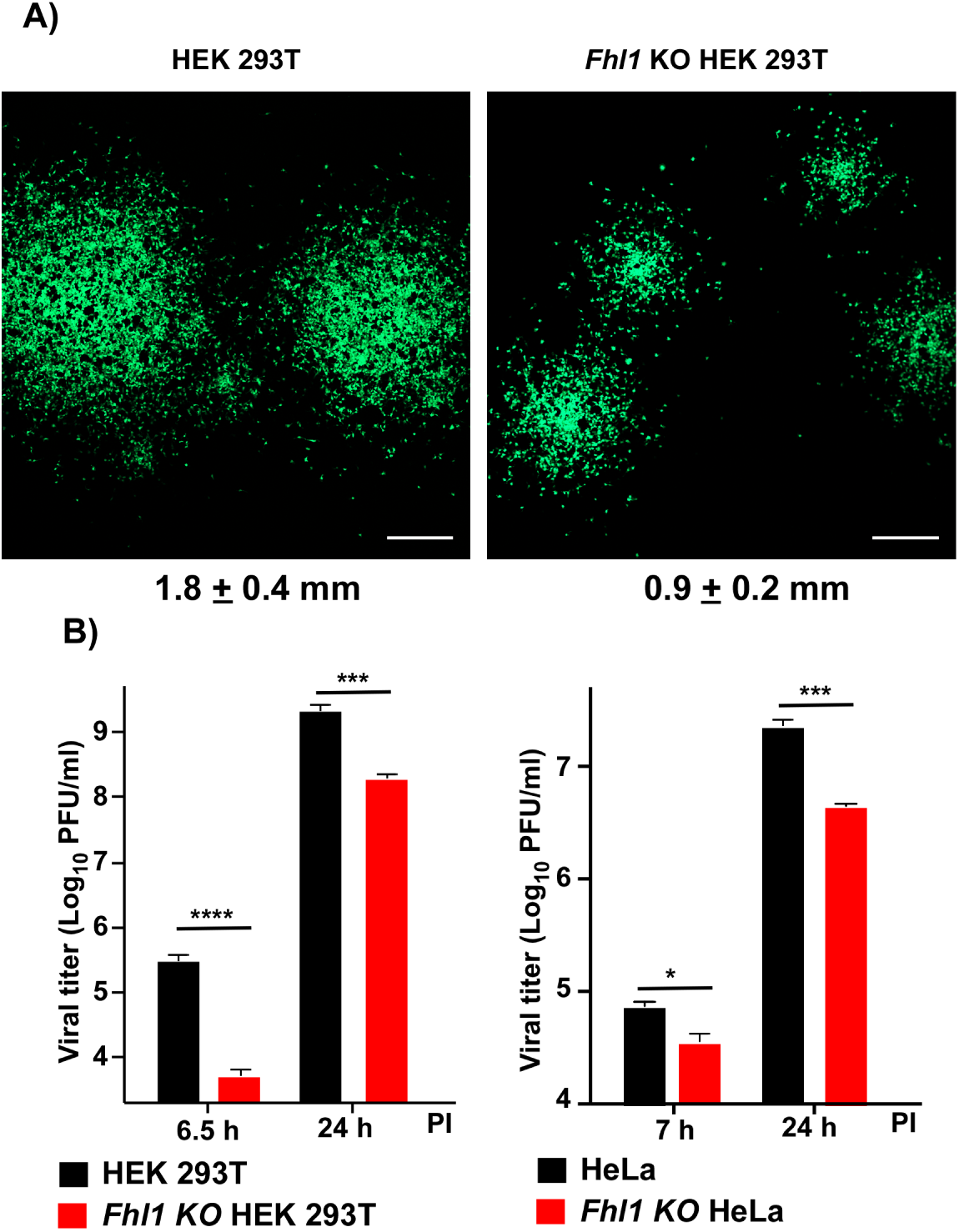
At low MOI, KO of *Fhl1* has a detectable negative effect on CHIKV/GFP titers. A) *Fhl1* KO HEK 293T and parental HEK 293T cells (2×10^6^ cells per well) were infected with CHIKV/GFP and incubated for 24 h at 37°C in Avicel-containing media as described in Materials and Methods. Then cells were fixed with 4% paraformaldehyde, and images of GFP-positive foci were acquired on a confocal microscope. Scale bar indicates 0.5 mm. Median diameters of GFP-positive foci and SDs are presented. The difference is statistically significant, p<0.001. B) *Fhl1* KO, and parental HEK 293T and HeLa cells were infected with CHIKV/GFP at an MOI of 0.1 PFU/cell. Viral titers were assessed at 6.5 and 24 h PI for HEK 293T and corresponding KO, and 7 and 24 h PI for HeLa and its KO. The experiment was repeated 3 times, means and SDs are indicated. Significance of differences were determined by two-way ANOVA Sidak test (*, P<0.05; **, P<0.01; ***, P<0.001; ****, P<0.0001; n=3).

To further dissect the effect of *Fhl1* KO, the parental HEK 293T and *Fhl1* KO HEK 293T cells were infected at an MOI of 0.1 PFU/cell, and viral titers were assessed at 6.5 and 24 h PI. At this low MOI, the differences in the rates of infectious virus release became detectable (Fig. 6B). At 6.5 h PI, the *Fhl1* KO cells reproducibly produced less virus and even at 24 h PI, viral titers remained significantly lower. At the latter time point, not all of the KO cells remained GFP-positive, but within the next 12 h, the infection reproducibly progressed to completion, and cells developed CPE. At the low MOI of 0.1 PFU/cell, at 7 and 24 h PI, CHIKV/GFP also accumulated to lower titers in the media of *Fhl1* KO HeLa cells, compared to the parental cell line (Fig. 6B).

The above data demonstrated that at least in some human cells, FHL1 has detectable pro-viral activity. Since CHIKV HVD functions during the assembly of viral replication complexes, it was reasonable to expect that lack of FHL1 expression in *Fhl1* KO HEK 293T cells may affect viral infectivity. To confirm this possibility, we titrated the stock of CHIKV/GFP at the same time on BHK-21, HEK 293T and *Fhl1* KO HEK293T cells. In repeat experiments, infectious titers determined on parental HEK 293T cells were indistinguishable from those determined on BHK-21. However, in the same experiments, titers determined on *Fhl1* KO HEK293T were reproducibly 6-8 times lower. Since the experiments were performed with the same virus, whose entry is dependent on heparan sulfate (38, 46) and on the derivatives of the same HEK 293T cells, this was an indication that KO of *Fhl1* has a negative impact on the ability of CHIKV/GFP to establish replication, but not on the viral entry.

To additionally confirm the detected pro-viral effects of FHL1 expression, the clone of *Fhl1* KO HEK293T cells, which was applied in the above experiments, was used to establish the stable cell line expressing FHL1-GFP fusion protein (*Fhl1* KO/KI HEK 293T). To reduce the possibility of nonspecific effects of ectopic overexpression, we selected for this study a cell clone with the lowest detectable level of GFP fluorescence, in which the level of protein expression was comparable to that in parental HEK 293T cells (Fig. 7A). In this KO/KI cells, replication of CHIKV/GFP was completely restored (Fig. 7B). In repeat experiments, both at early and late times PI, titers of viruses harvested from the original HEK 293T cells and its *Fhl1* KO/KI derivative were identical at the used MOI.

**FIG. 7.**
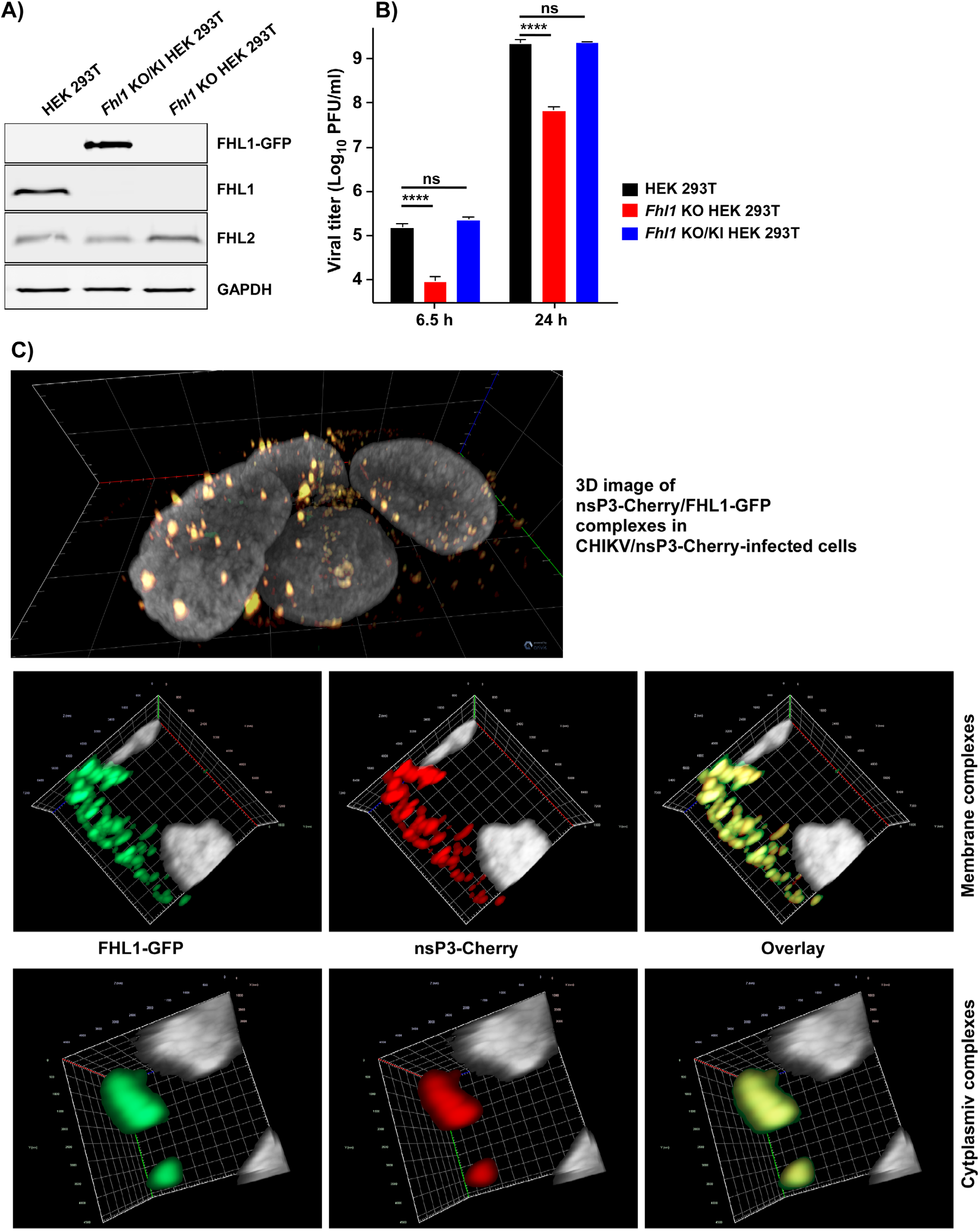
Expression of hFHL1-GFP fusion in *Fhl1* KO HEK 293T cells restores replication of CHIKV/GFP to the wt level. A) WB analysis of FHL1 and FHL2 expression in the parental cells and generated KO/KI cell line. The membranes were stained with FHL1- and FHL2-specific Abs and scanned on LI-COR imager as described in Materials and Methods. B) The indicated cell lines were infected at an MOI of 0.1 PFU/cell with CHIKV/GFP. At the indicated time points, viral titers were determined by plaque assay on BHK-21 cells. C) *Fhl1* KO/KI cells were infected with CHIKV/nsP3- Cherry at an MOI of 1 PFU/cell. At 7 h PI, cells were fixed and counterstained for nuclei with DAPI. 3D images were acquired on Zeiss LSM800 confocal microscope in Airyscan mode and processed and assembled in ZEN 2.6 (Blue edition) software.

The expressed GFP-FHL1 fusion was diffusely distributed in the uninfected cells. However, within 6 h PI with CHIKV/nsP3-Cherry, which encodes fluorescent Cherry protein in nsP3 HVD, GFP-FHL1 concentrated in both membrane-bound pre-vRCs and large cytoplasmic complexes formed by nsP3-Cherry, indicating an interaction between these two proteins (Fig. 7C).

Taken together, the above data demonstrated that the expression of FHL1 has a stimulatory effect on CHIKV infection in human cells, but the pro-viral effect is not as strong as that previously described for G3BPs. Thus, replication of CHIKV/GFP can proceed in the absence of FHL1 expression, albeit the early step(s) of viral replication are detectably affected.

### Binding of FHL1 to CHIKV HVD is mediated by the LIM1 domain

We have previously assigned the backbone resonances in the NMR spectra of the entire CHIKV HVD (25). In this new study, these data allowed us to use the NMR titration approach, which is based on observation of the changes in ^1^H-^15^N best-Trosy spectra, to precisely identify the binding site of FHL1. Upon addition of unlabeled FHL1 to uniformly ^15^N-labeled CHIKV HVD (Fig. 8A), we observed gradual decreases of intensities of cross-peaks of amide backbone resonances (Fig. 8B) in two HVD fragments. The cross-peak intensities for the first fragment (aa 412-458) were broadening with the increase in concentration of FHL1 and completely disappeared at the ratios of HVD to FHL1 above 1:0.6. No new cross-peaks were found at 1:1.2 ratio. It is a common effect that cross-peaks in disordered peptide completely disappears after ligand binding (47, 48), but the exact mechanism of this phenomenon for the intrinsically disordered proteins (IDPs) remains unknown.

**FIG. 8.**
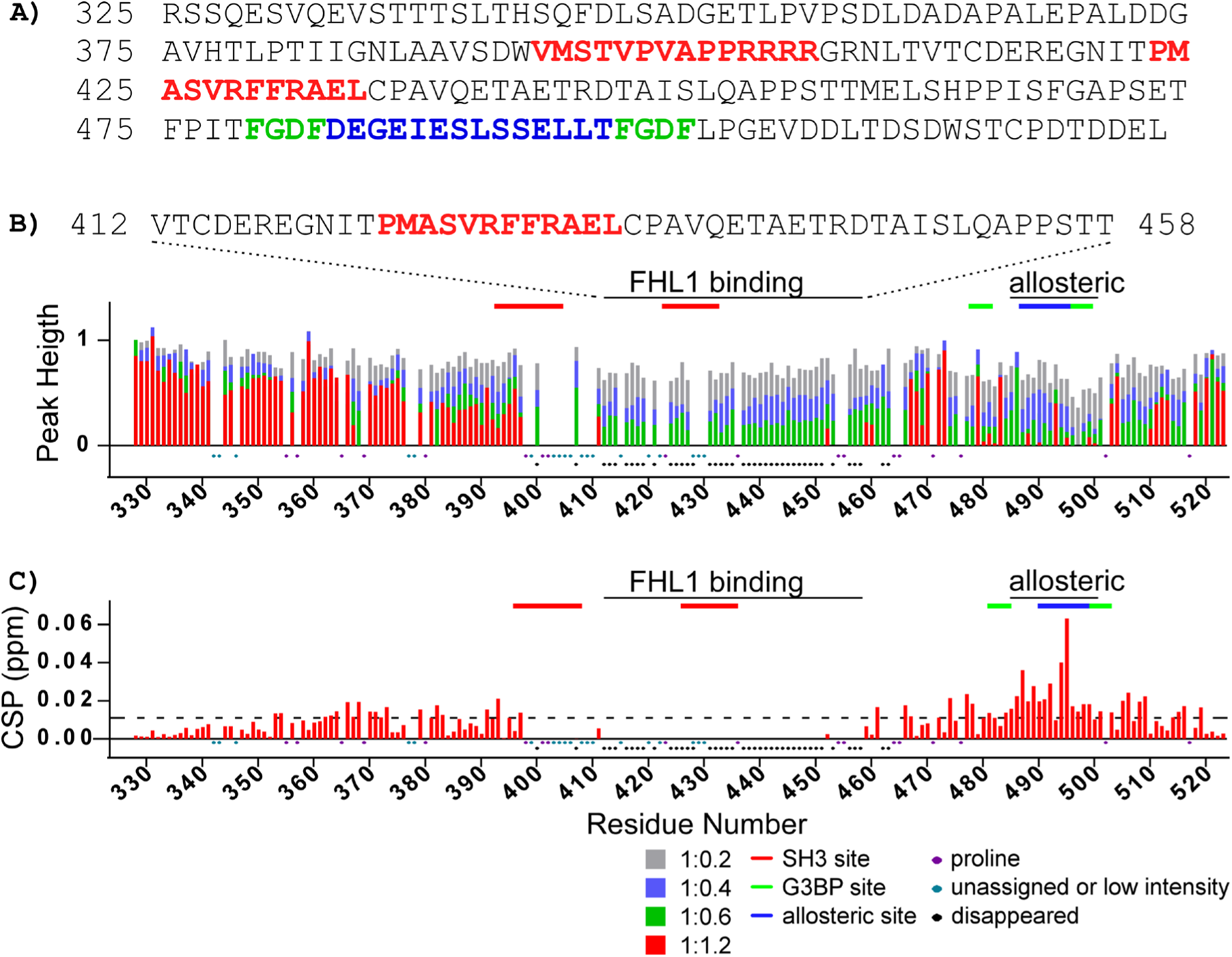
Changes of the CHIKV HVD amide resonances in ^1^H-^15^N best-TROSY spectra upon titration with FHL1. A) The aa sequence of CHIKV nsP3 HVD, which was used in the experiments. The previously identified CD2AP- and G3BP-binding, and allosteric sites are indicated in red, green and blue colors, respectively. Both the changes in normalized intensities (Peak height) as well as chemical shifts (CSP) are shown in panels (B) and (C), respectively. In panel (C) the dashed line shows the level of 2 standard deviations of variation of CS during titration. Ratio of titrations used are shown below panel (C). Prolines, unassigned aa, low intensities or cross-peaks disappearing during titration are indicated below panel (C). Marked above in red solid lines are the CD2AP-binding sites, in blue-the proposed CD2AP-specific allosteric site and in green-the G3BP-binding sites. They are marked correspondingly to the aa sequence in (A).

The identified HVD fragment strongly overlaps with the previously described (25) SH3-binding site 2 (marked by the red line in Fig. 8B). We excluded the SH3 domain-binding site 1 from the analysis since even in the apo form, the non-proline cross-peaks in this region (aa 399-410) either had low intensities or strongly overlapped with other cross-peaks. This precluded correct quantification of their intensities. Based on the data, we concluded that the peptide between aa 411-459 represents a possible FHL1-binding site. It starts from the aa sequence located between two SH3 domain-binding sites of CD2AP and extends well beyond the second SH3 binding site. It is not uncommon for LIM domain-containing protein to occupy large binding sites due to extended interactions of disordered peptides with the LIM domain surface (49, 50).

The cross-peaks in another HVD fragment (aa 485-500) demonstrated different behavior. Noteworthy, this fragment covers the previously proposed G3BP-binding site (19). Their intensities were decreasing with the increase in concentration of FHL1 up till ratio 1:0.6 and new cross-peaks were readily detectable (Fig. 8C) with increasing intensity till ratio 1:1.2. This type of behavior indicates that the system is in slow exchange regime (51). Interestingly, we have previously identified a possible allosteric site in this C-terminal HVD fragment, which was responsive to binding of individual SH3 domains of CD2AP and their natural combination (SH3-All) (25). To confirm that this distant fragment behaves similarly upon binding either FHL1 or CD2AP, we compared the chemical shift perturbation (CSP) trajectories specific for FHL1 and CD2AP SH3-B domain (25) complexes (Fig. 10). The trajectories of the individual amides, even though they are unique, were identical irrespective of FHL1 or SH3-B domain binding. Thus, the aa 485-500 fragment of HVD represents a possible allosteric site. The nature of the allostery is currently under investigation.

Next, we attempted to further dissect the FHL1 binding to CHIKV HVD by analyzing HVD interactions with individual LIM domains. Note that the LIM1 construct included the N-terminal part of FHL1, which contained natural half-LIM domain. We did not find significant reductions in cross-peak intensities even at 1:5 ratio of HVD to LIM2 (Fig. 9). This suggested that LIM2 alone does not interact with CHIKV HVD. However, three other LIM domains demonstrated detectable bindings with LIM1-HVD interaction being the strongest. The intensities of the majority cross-peaks in the binding fragment for LIM1-HVD complex were already reduced at a 1:1 ratio and completely disappeared at 1:5 ratio. Similar to what was observed for the entire FHL1, no new cross-peaks were detected. Interestingly the LIM1-binding site was shorter than that of the entire FHL1, but it still remained 36-aa-long (aa 422-457). We also observed reduction in cross-peak intensities in the allosteric site. However, these cross-peaks were in fast exchange, and the corresponding transient chemical shifts were readily detectable (Fig. 10). Thus, LIM1 (with the N-terminal half-LIM domain) appears to be mainly responsible for FHL1 binding to CHIKV HVD, but its binding does not exactly mirror that of the entire FHL1.

**FIG. 9.**
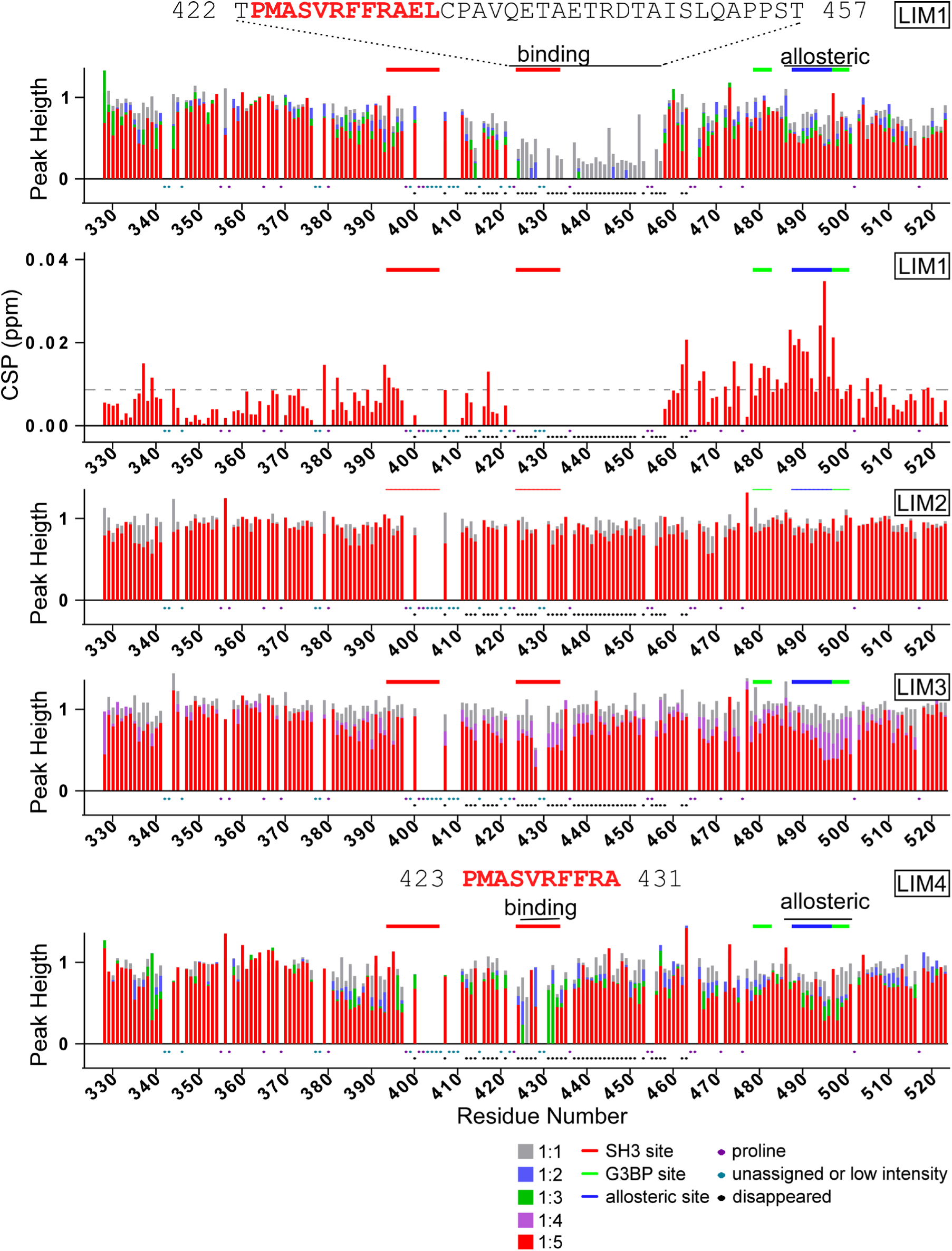
Changes of the CHIKV HVD amide resonances in ^1^H-^15^N best-TROSY spectra upon titration with LIM1, LIM2, LIM3 and LIM4 domains. For the titration with LIM1, both the changes in normalized intensities (Peak height) as well as chemical shifts (CSP) are shown (two panels from the top, respectively). For LIM2, LIM3 and LIM4 only the changes in normalized intensities (Peak height) are shown. The level of 2 standard deviations of variation of CS during titration is shown by dashed line. Ratio of titrations used are shown below panels. Prolines, unassigned aa, low intensities or cross-peaks disappearing during titration are indicate below panels. Marked above in red solid lines are the CD2AP-binding sites, in blue-the proposed CD2AP-specific allosteric site and in green-the G3BP-binding sites.

**FIG. 10.**
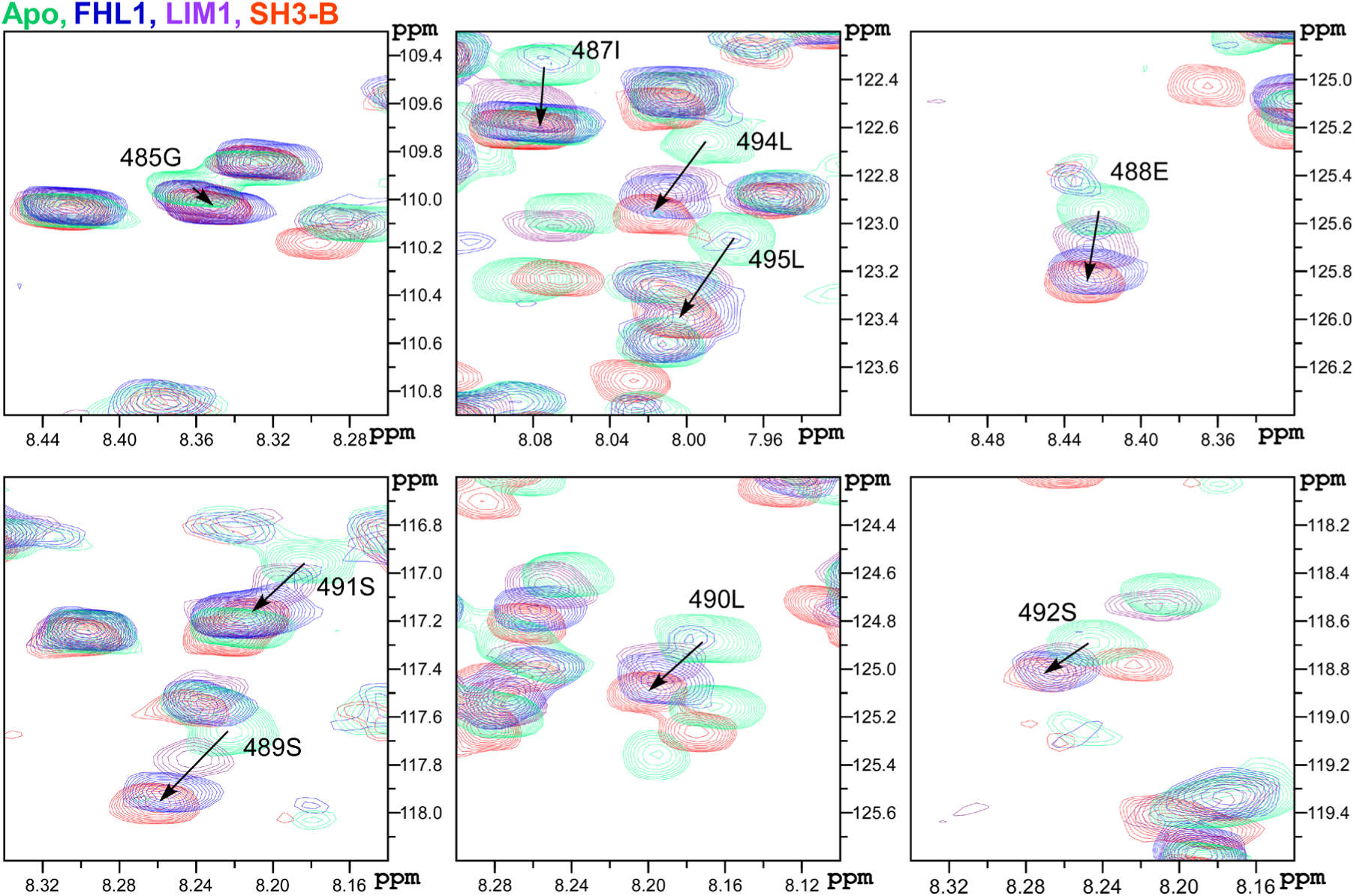
Chemical shift perturbation signature of HVD binding with different ligands at the possible allosteric site. Expanded ^1^H-^15^N best-TROSY spectra for the aa belonging to the possible allosteric site of CHIKV HVD. The ^15^N-labelled CHIKV HVD (coloured by green) was mixed with unlabelled FHL1 (1:1.2 ratio) and LIM1 (1:5 ratio) coloured by blue and purple respectively. Data for SH3-B (1:5 ratio) (red) were taken from previous publication (25). The chosen ratio represents the maximum CSP observed under titration. The shift trajectories shown by arrows for all ligands for aa in this site are similar.

Two other domains, LIM3 and LIM4, demonstrated essentially weaker interactions with HVD. Nevertheless, the intensities of the cross-peaks of the aa 423-431 fragment were reduced or completely disappeared with the increase in concentration of LIM4 (Fig. 9). The latter site includes the previously identified binding site 2 of CD2AP (25). We also observed a slight reduction of intensity for this HVD fragment in titration experiments with LIM3.

Interestingly, the HVD fragment of aa 423-431, which interacts with three LIM domains, contained two adjacent phenylalanines (aa 429 and 430), whose amide resonances were not observed in the spectra of CHIKV HVD, most likely due to some sort of interaction, possibly stacking. We hypothesized that LIM1, LIM3 and LIM4 domains may contain similar hydrophobic binding pockets that bind one of these phenylalanines or both. Upon evaluation of the 3D structures of different LIM domains, we identified in all of them the apolar pockets formed by four to five aromatic residues, which include a tryptophan, one or two tyrosines and two or three phenylalanines. In LIM1 domain, the proposed apolar pocket is formed by Y56, W61, F66 and F80 (Fig. 11A), and is localized in a groove and stabilized by two β-strands. We speculate that binding of LIM1 to HVD could promote the displacement of F80 and insertion of one of the above-described HVD-specific phenylalanines into the indicated apolar pocket. Good alignment was found between the apolar pockets of LIM1 and LIM4 (Fig. 11B), with the minor difference in the presence of additional phenylalanine in LIM4 pocket. Alignment of LIM1 structure with LIM2 and LIM3 revealed larger differences, particularly for the LIM2 domain. The apolar pocket in LIM2 and LIM3 is stabilized by two pairs of β-strands, one pair on each side of the pocket, which possibly complicate interaction with F429F430 of HVD. These differences could explain the lower binding affinity of the LIM3 domain and the absence of interaction of LIM2 domain with CHIKV HVD. Further experimental proof for the critical role of F429F430 in CHIKV HVD in the FHL1 binding is presented in the following section.

**FIG. 11.**
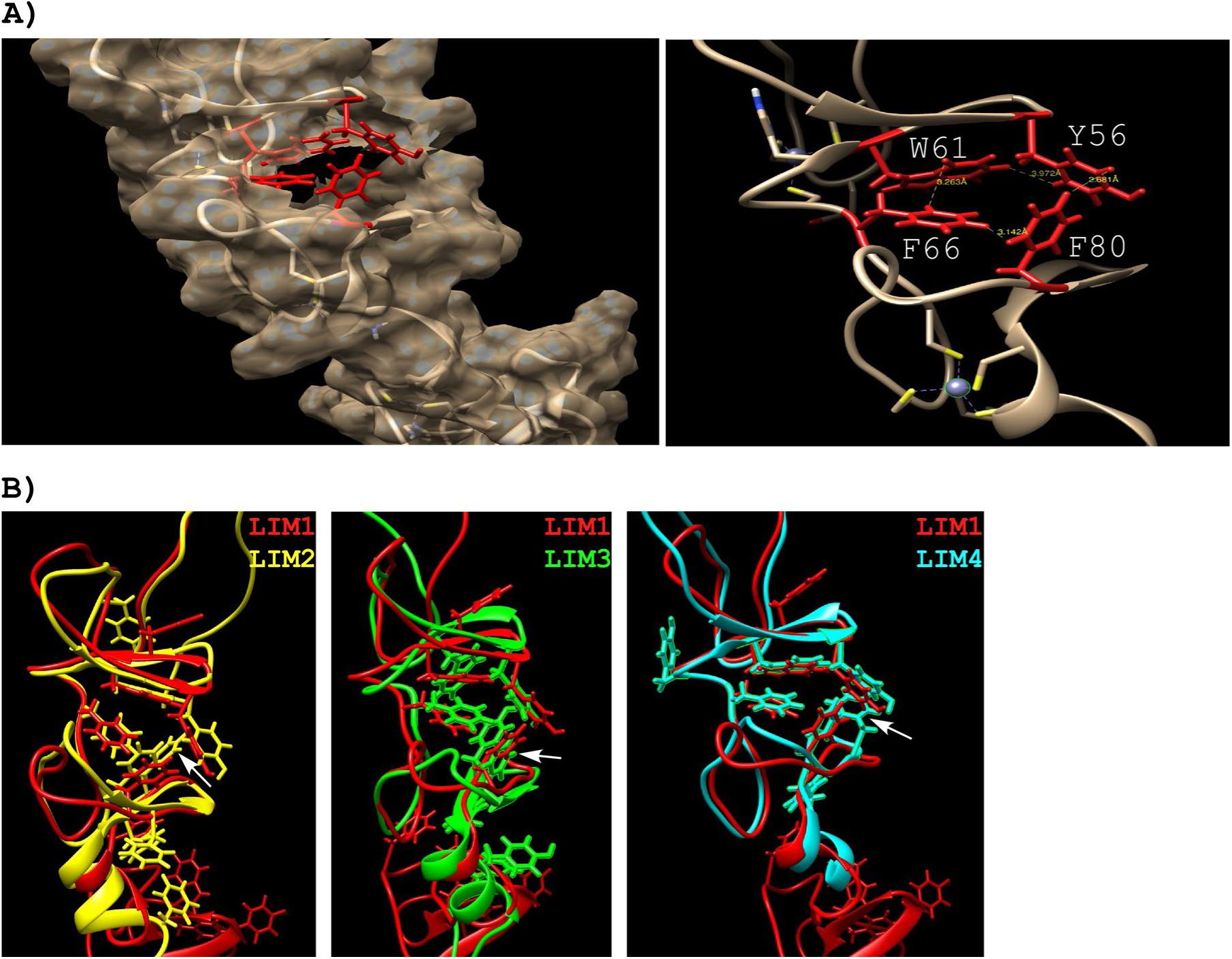
Structures of the individual LIM domains. A) LIM1 (PDB 2cup) showing the surface except around the apolar site (left panel), where the sidechains are in red: Y56, W61, F66 and F80. The expansion (right panel) shows the close proximity of the aromatic residues in the apolar site. (B) Pairwise comparison between LIM1 (PDB 2cup, red) and LIM2 (PDB 1×63, yellow), LIM3 (PDB 2cur, green) and LIM4 (PDB 2egq, cyan).

The higher affinity of the LIM1 domain could also be determined by additional interactions of the HVD-binding site with the surface of LIM1 domain. The alignment presented in Fig. 12 includes all non-redundant CHIKV sequences, which are currently available in GeneBank. Two fragments with higher conservation, which could potentially determine such interactions, are evident. The first one is located between two CD2AP-binding sites and is likely involved in binding of the full-length FHL1. The second, downstream-located fragment, extends over 11 aa and does not overlap with the binding site of CD2AP. Both of these sequences are located in the FHL1-binding site determined in the above NMR study.

**FIG. 12.**
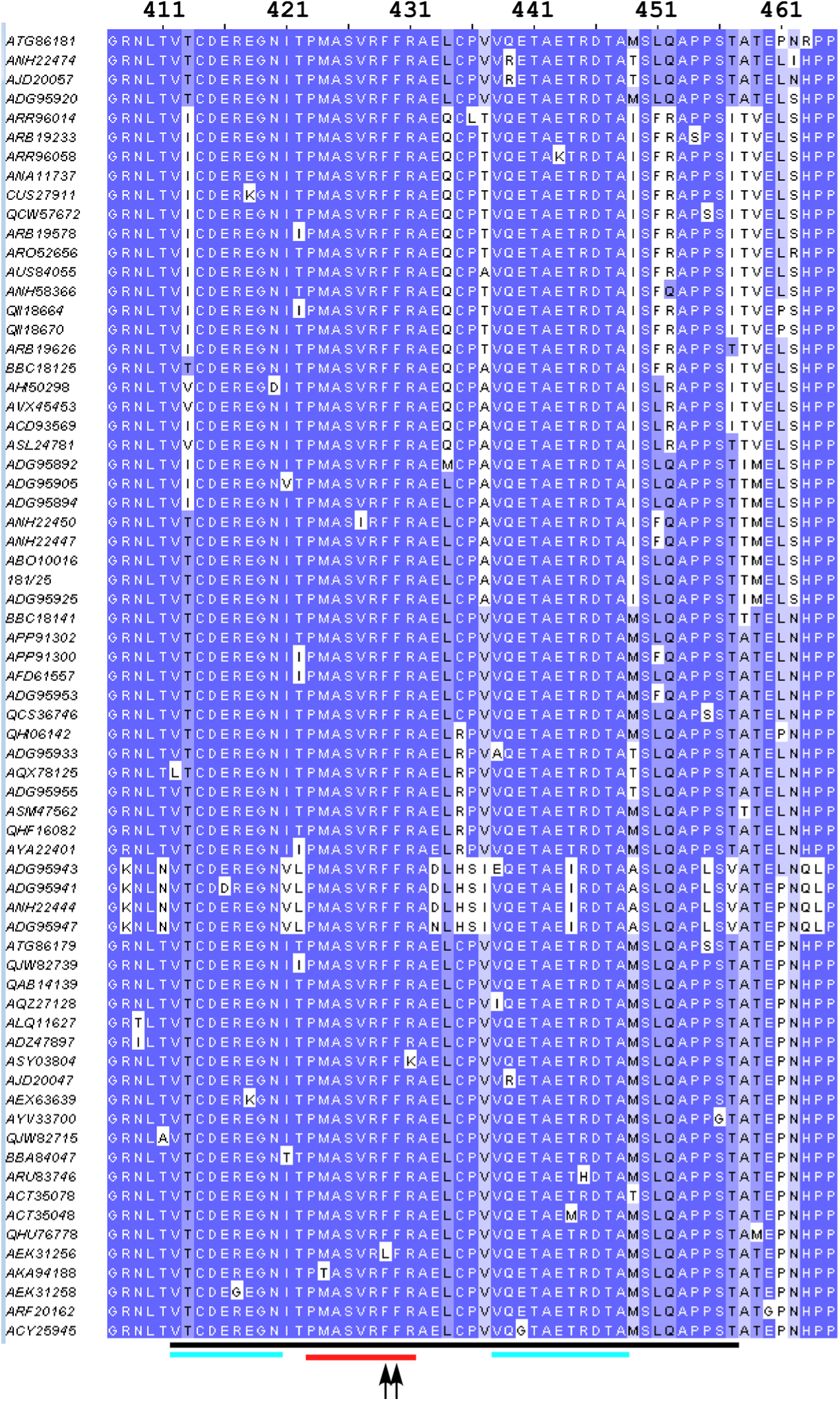
Alignment of CHIKV nsP3 HVD fragment that contains FHL1-binding site. The aa sequences of the FHL1-binding fragment of CHIKV sequences in GenBank were downloaded from GeneBank and aligned using Jalview 2.11.10. The redundant sequences were deleted. The black line indicates the predicted FHL1-binding site. The blue lines indicate the aa sequences with highest levels of identity. The red line indicates the second binding site of CD2AP. The black arrows point to F429 and F430.

The NMR data demonstrated overlap between the FHL1- and CD2AP-binding sites in CHIKV HVD and suggested a possibility of competition between the two proteins for binding to HVD. To experimentally evaluate this possibility, we purified the full-length FHL1 and analyzed its binding with CHIKV HVD by gel exclusion chromatography. We found that FHL1 directly binds to CHIKV HVD and forms a stable dimeric complex (Fig. 13A). In agreement with our previous data (25), the N-terminal fragment of CD2AP (SH3-All), which contains all SH3 domains, also formed a stable dimeric complex with CHIKV HVD (Fig. 13B). Next, we tested whether FHL1 and SH3-All can simultaneously bind to CHIKV HVD. Indeed, a trimeric complex containing CHIKV HVD, FHL1 and SH3-All was clearly present (Fig. 13C). Thus, even if binding sites for FHL1 and CD2AP overlap, these two proteins do not appear to compete for binding to CHIKV HVD. We have also confirmed that FHL1 and SH3-All do not directly interact (Fig. 13D). Although, this does not rule out a possibility that the C-terminal disordered domain of CD2AP, which is omitted in our construct, can potentially mediate an additional interaction with FHL1.

**FIG. 13.**
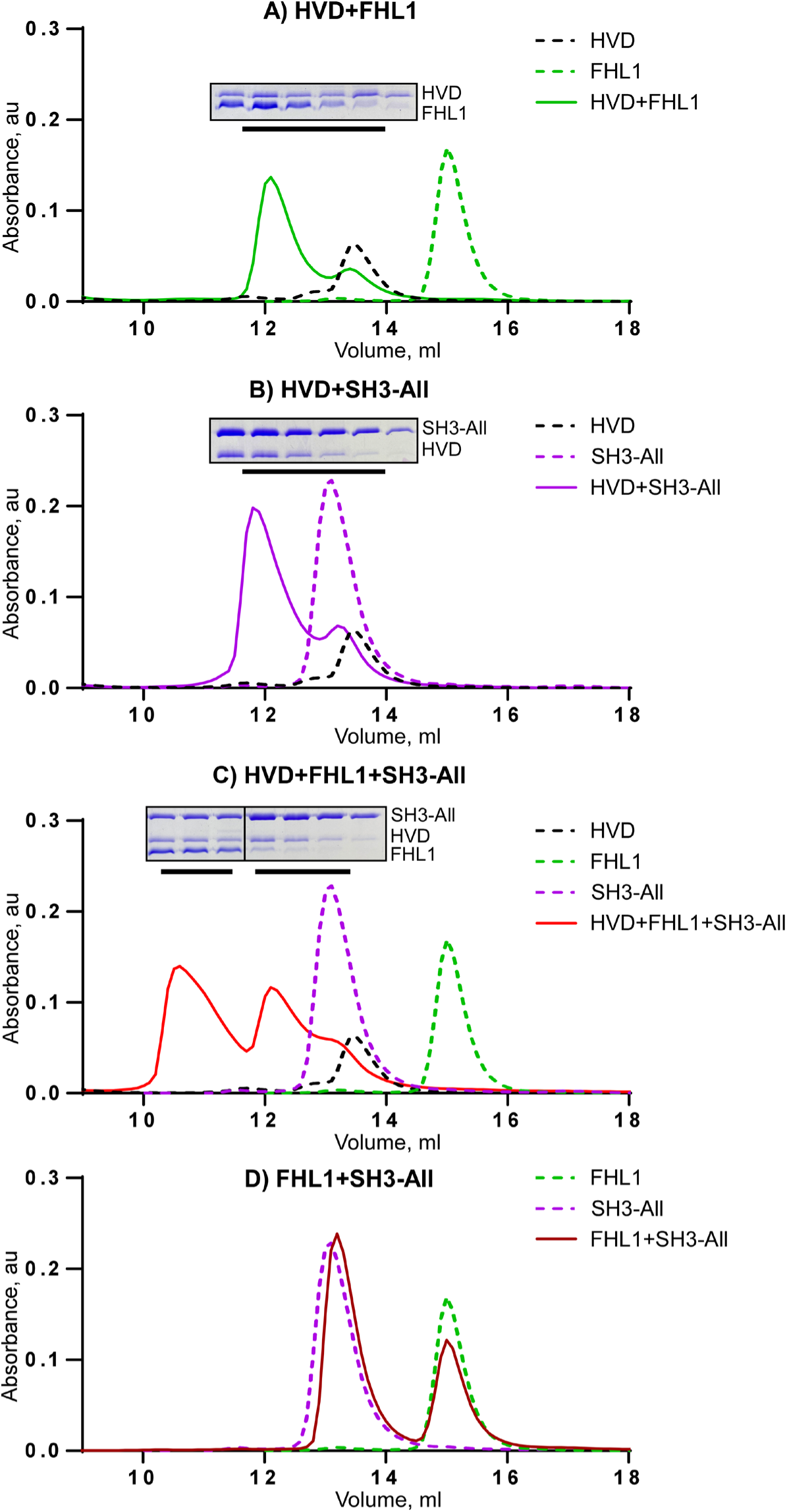
CHIKV HVD, FHL1 and SH3-All form trimeric complex in solution. Analyses of binding were performed by size exclusion chromatography on Superdex^TM^ 200 increase 10/30 GL column. A) CHIKV HVD + FHL1, B) CHIKV HVD + SH3-All, C) CHIKV HVD + FHL1 + SH3-All, D) FHL1 + SH3-All. Fraction corresponding to the peaks indicated by black lines were analyzed by SDS-PAGE. Gels were stained with Coomassie.

In conclusion, i) we have identified a binding site for FHL1 in CHIKV HVD and demonstrated that the FHL1-HVD interaction is primarily mediated by LIM1 and the N-terminal half of LIM domains. We hypothesize that ii) FHL1 binding is determined by the interaction of F429F430 in HVD with the apolar pocket of LIM1 and by other conserved aa sequences in CHIKV nsP3 HVD with the LIM1 surface. iii) Binding of FHL1 induces allosteric changes in the distant, C-terminal fragment of CHIKV HVD. iv) Despite having overlapping binding sites, CD2AP and FHL1 interact independently with CHIKV HVD and form a trimeric CD2AP/HVD/FHL1 complex. Further structural studies are needed to resolve this complex structure.

### Mutations in the FHL1-binding site of nsP3 HVD attenuate CHIKV replication

To further define the FHL1-binding site and the biological significance of the structural data, we designed a wide range of CHIKV/GFP variants with mutations in the HVD (Fig.14A). They contained combinations of mutations in the predicted FHL1-binding site (M1, M2, M3, M6, M7, M8) or in both CD2AP- and FHL1-binding sites (M4, M5 and M9). The *in vitro*-synthesized RNAs of the mutants were electroporated into BHK-21 cells, and in the infectious center assay, they demonstrated infectivities similar to that of parental CHIKV/GFP RNA. This was an indication that no additional adaptive mutations were required for viral viability in BHK-21 cells. To avoid possible viral evolution, the generated stocks were used for further experiments without additional passaging.

**FIG. 14.**
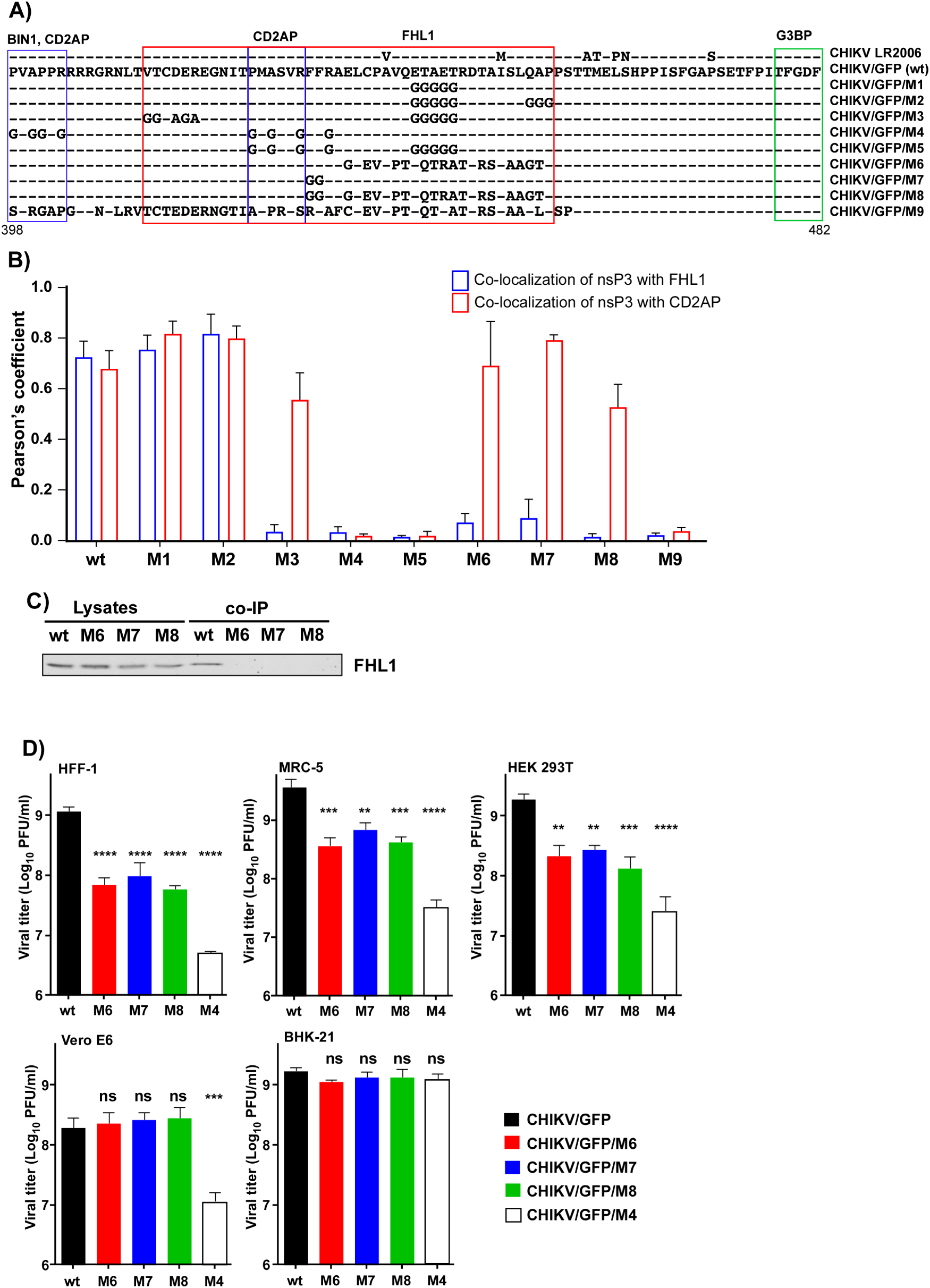
Inactivation of FHL1- and CD2AP-binding sites has additive negative effects on CHIKV/GFP replication but does not make virus nonviable. A) Alignment of the aa sequences of CHIKV nsP3 HVD fragment of CHIKV 181/25 and generated HVD mutants. Binding sites for FHL1 and CD2AP, which were identified in the above-described and previous NMR studies, are indicated by red and blue boxes, respectively, and G3BP-binding repeat is indicated by green box. B) HEK 293T cells were infected with the indicated CHIKV variants at an MOI of 1 PFU/cell. At 15 h PI, cells were fixed and stained with antibodies specific to CHIKV nsP3 and FHL1 or CD2AP and secondary Abs. Pearson’s coefficients (means and SDs are shown). C) HEK 293T cells were infected with VEEV replicons expressing Flag-GFP fused with HVDs of the indicated mutants. HVD complexes were isolated using beads with Flag-specific MAbs and analyzed by WB using FHL1-specific Abs. D) The indicated cell lines were infected with the parental CHIKV/GFP and its nsP3 HVD mutants at an MOI of 0.1 PFU/cell. Viral titers were determined at 24 h PI by plaque assay on BHK-21 cells. The experiment was repeated 3 times, means and SDs are indicated. Significance of differences were determined by one-way ANOVA Dunnett test (*, P<0.05; **, P<0.01; ***, P<0.001; ****, P<0.0001).

First, we evaluated the ability of nsP3 of the designed mutants to make complexes with FHL1 and CD2AP. HEK 293T cells were infected with all mutants and the parental CHIKV/GFP. They were fixed at 15 h PI and stained with antibodies specific to CHIKV nsP3 and FHL1 or CD2AP. Multiple random images were taken for each mutant and the levels of nsP3/FHL1 and nsP3/CD2AP co-localization were assessed based on Pearson’s coefficient (Fig.14B). Co-localizations of nsP3 with either FHL1 or CD2AP were unaffected by the mutations in M1 and M2. However, randomization of a 21-aa-long peptide downstream of CD2AP-binding site made M6 mutant incapable of interacting with FHL1, but not with CD2AP. The same effect was observed with M3 mutant that contained mutations in the conserved fragments located upstream and downstream of the second CD2AP-binding site. Mutation of F429F430 alone or in combination with 21-aa-long mutation also strongly affected the ability of M7 and M8 mutants, respectively, to bind FHL1, while interaction with CD2AP was not affected. M4, M5 and M9 mutants contained mutations in the previously described CD2AP-binding site (aa 423-428) and were unable to make complexes with either FHL1 or CD2AP.

To confirm the effect of the 21-aa-long and F429F430 mutations on FHL1-HVD interaction, the mutated and wt HVDs were expressed from VEEV replicon as fusions with Flag-GFP (see Materials and methods for details). CHIKV HVD complexes were isolated using Flag-specific Abs and analyzed by WB using FHL1-specific Abs. No FHL1 was detected in the co-IP samples of M6-, M7- and M8-derived HVD (Fig. 14C).

Taken together, the IF and co-IP data confirmed the NMR results. i) The FHL1-interacting peptide of HVD is unusually long. ii) It overlaps with the second binding site of CD2AP and includes the peptides located upstream and downstream of the latter site. iii) F429F430 di-peptide, which was proposed to interact with the apolar pocket in LIM1 domain, likely plays a critical role in this interaction, which is comparable to the effects of longer aa sequences.

Next, BHK-21, MRC-5, HFF-1, HEK 293T and Vero cells were infected at an MOI of 0.1 PFU/cell with i) M6, M7 and M8 mutants, encoding HVDs unable to interact with FHL1, ii) M4 mutant, whose HVD does not interact with either FHL1 or CD2AP, and iii) the parental CHIKV/GFP. Viral titers were evaluated at 24 h PI (Fig. 14D). BHK-21 cells were used as a highly susceptible cell line for CHIKV infection, and MRC- 5 and HFF-1 cells were applied because they are close to primary human fibroblasts that are believed to be a main primary target of CHIKV infection *in vivo* (52).

In repeat experiments, the effects of mutations in the FHL1-binding site on replication of all four mutants in BHK-21 cells were undetectable. Since BHK-21 cells have previously been found to support replication of alphaviruses with strongly modified genomes (15, 19, 53), this result was not surprising. Alterations of replication of the designed HVD mutants were more prominent in MRC-5, HFF-1 and HEK 293T cells. At 24 h PI, M6, M7 and M8 mutants demonstrated ∼10-fold lower titers, and the infectious titers of the released M4 mutant were reduced by 2 orders of magnitude. However, all of the mutants remained capable of developing spreading infection. Interestingly, in Vero cells, titers of M6, M7 and M8 mutants were not significantly different from those of parental CHIKV/GFP, and only replication of M4 was affected by introduced mutations. These data correlate with the undetectable level of FHL1 in Vero cells (Fig. 1A), resulting in an inability of the HVD-FHL1 to play a role in supporting replication of the virus with wt HVD. Taken together, the data were in agreement with the results of the above-described experiments with *Fhl1* KO cells. They supported the hypothesis that interactions of FHL1 and SH3 domain-containing proteins, such as CD2AP, with CHIKV nsP3 HVD have additive positive effects on viral replication and infectious titers, but are generally dispensable. CHIKV mutants encoding HVD incapable of interacting even with both groups of proteins remain viable.

## DISCUSSION

Replication of CHIKV and other alphaviruses is critically determined by the host factors that mediate the assembly of vRCs and function in recruitment of viral genome and synthesis of both G and SG RNAs. CHIKV nsP3 and its HVD in particular play indispensable roles in these processes. This long, intrinsically disordered HVD contains a combination of short motifs that interact with cell-specific sets of host factors and functions as a hub for vRC assembly. The previous studies identified a group of the SH3 domain-containing proteins along with G3BP, FHL and NAPL1 family members as CHIKV HVD-interacting partners that have the abundance in the nsP3 complexes (16, 17, 19, 20). However, the mechanisms of their functions remain to be fully understood. The currently available data can be summarized as follows. I) CHIKV replication does not rely on single members of the host protein families but is rather redundantly determined by all of the family members or even by the representatives of different families having similar domains, such as the SH3 domain-containing proteins. II) CHIKV HVD interactions with G3BP family members are indispensable for viral RNA replication, and their alterations make virus nonviable. III) Extensive mutagenesis of HVD, which prevents binding of all of the host factors except G3BPs, also abolishes CHIKV replication. IV) Restoration of at least one of the binding motifs in CHIKV HVD in addition to those interacting with G3BPs makes such variants viable but replicate less efficiently than the wt virus. These data suggested a high level of redundancy in the functions of different families of host factors and their additive activities in CHIKV RNA replication. However, this redundancy also complicates the dissection of binding sites and the roles of particular host proteins in vRC formation and function.

Previously, by using structural, biochemical and biological approaches, we and other groups have characterized the interactions of G3BP1/2, and CD2AP, BIN1 and SH3KBP1 with CHIKV HVD (19, 20, 27, 28). In this new study, we were mostly focused on the analysis of FHL1-HVD interactions. FHL1 and FHL2 representatives of the FHL family contain four and a half LIM domains that demonstrate a strong level of conservation and have two zinc fingers (31). FHL1A isoform that was used in this study for ectopic expression is highly expressed in skeletal muscles. FHL2 is expressed in a wide range of tissues, such as bone marrow-derived dendritic and skeletal muscle cells, and exhibits affinities to various intracellular proteins and receptors (31). In the previous studies, we identified FHL1 or FHL2 in CHIKV HVD complexes isolated from human and mouse cells, respectively, but not both of them simultaneously. This was suggestive of their function(s) in vRC formation in a cell type-specific mode. Indeed, analysis of a variety of cell lines in terms of FHL1 and FHL2 expression demonstrated that both proteins are not ubiquitously expressed, and some human and mouse cells do not express FHL1 at all (Fig. 1). Despite such characteristics, these cells efficiently supported CHIKV replication suggesting that the latter protein is not an absolute pre-requisite of vRCs’ formation and function. In HEK 293 and HeLa cells, the KO of the *Fhl1* gene also had a detectable but relatively small negative effect on CHIKV growth that cannot be compared to the previously described deleterious effect of double KO of *G3bp1* and *G3bp2* (16, 29). Interestingly, the lack of expression of both FHL1 and FHL2 in mouse *Fhl1* KO NIH 3T3 cells failed to make these cells non permissive for CHIKV replication (Fig. 4). This was a strong indication that at least in these mouse cells, other host proteins can replace FHLs in vRCs.

Analysis of the FHL1-binding sites in CHIKV HVD by NMR spectrometry led to unexpected results. The FHL1-interacting site was found to be longer than previously studied binding motifs in alphavirus HVDs and partially overlap with one of the previously identified binding site of CD2AP (19, 20, 25). However, in biochemical assays, CD2AP and FHL1 did not demonstrate competition for binding peptide and were able to generate a triple HVD/CD2AP/FHL1 complex. On the other hand, we did not detect cooperativity in CD2AP and FHL1 binding to CHIKV HVD (Fig. 13). Viral mutants, which lost an ability to interact with FHL1, also remained capable of forming CD2AP/nsP3 complexes and were viable (Fig. 14). To date, the mechanism of formation of triple HVD/CD2AP/FHL1 complex remains to be better understood, but we speculate that CD2AP and FHL1 may interact with different amino acids in the same peptide in CHIKV HVD. Thus, these two host factors have rather additive than synergistic pro-viral effects.

As described above, FHL1 has four and a half LIM domains, which demonstrate high levels of conservation (31). However, in the NMR-based experiments only domain 1 was found to efficiently interact with CHIKV HVD. Binding of domains 3 and 4 was very inefficient, and domain 2 demonstrated no binding at all. Interestingly, the entire protein that combines all 4 LIM domains was found to interact with a wider fragment of HVD, suggesting that single domain-based data do not always fully reproduce the results obtained with larger proteins. A similar conclusion was made in our previous study when the interaction of the SH3-binding domains did not reproduce interaction of the entire CD2AP protein with CHIKV HVD (25).

Overlap of the CD2AP- and FHL1-binding sites complicated the dissection of the effects of interaction of each protein with HVD on viral replication. Despite our efforts, we were unable to design an HVD mutant that could interact with FHL1 without CD2AP binding. However, inactivation of the FHL binding while saving HVD interaction with CD2AP was possible. In agreement with the results from the experiments with *Fhl1* KO cells, CHIKV mutants, which no longer formed FHL1-HVD complexes, demonstrated a significant decrease in replication. Nevertheless, they remained viable in the used vertebrate cell lines, formed nsP3/CD2AP complexes in the infected cells and were capable of developing a spreading infection. Interestingly, mutation of both conserved phenylalanines (F429F430) had a strong negative effect on CHIKV infection that was similar to the effect caused by mutation of a 21-aa-long downstream-located peptide. We speculate that this FF di-peptide may play a critical role in FHL1-HVD binding by interacting with hydrophobic pocket formed in FHL1 LIM domains. However, this hypothesis needs to be confirmed by further structural work on atomic resolution.

CHIKV mutant with mutations in HVD that made it unable to make complexes with both FHL1 and CD2AP showed less efficient replication than those having mutated FHL1-binding site only. Thus, the interaction of CHIKV with FHL1 and CD2AP likely has an additive pro-viral effect, but the elimination of even both interactions is not lethal for the virus. This conclusion is supported by our previous data and the results from the studies of other groups (19, 20, 25). At that time, there was no structural information that CD2AP- and FHL1-binding sites overlap. Thus, the effects of the introduced mutations were interpreted as alterations of CD2AP binding. However, according to our new findings, it was more complex, and mutations likely abolished FHL1-HVD interaction as well.

In recent studies, we have already successfully applied our accumulating structural and functional knowledge about HVDs for the development of the next generation of vaccine candidates of EEEV and VEEV infections (54, 55). The extensive mutations or deletions of the host protein-binding motifs in their HVDs made such viral mutants attenuated *in vivo* while remaining highly immunogenic. Thus, extensive modifications in the alphavirus HVDs greatly improve the safety and stability of attenuated phenotypes. In case of CHIKV, they may be applied either alone or in combination with i) the previously selected mutations in E2 glycoprotein (38), ii) with the recently identified cluster mutations in the V peptide in nsP2, which make the latter protein incapable of interfering with the development of the innate immune response (45), and/or iii) with mutations in the macro domain of nsP3, which affect the ability of CHIKV to inhibit cellular translation (56).

In conclusion, the results of this study demonstrate that i) the interaction(s) between members of the FHL family of cellular proteins and CHIKV nsP3 has pro-viral functions. As we have previously described for the SH3 domain-containing proteins, such as CD2AP, this FHL-HVD interaction plays a significant role in early stages of CHIKV replication and thus, increases viral infectivity. However, it is not a pre-requisite for viral replication, and CHIKV is capable of developing spreading infection in the *Fhl1* KO cells and the cell lines that naturally do not express FHL1. ii) FHL1-binding site is located between aa 411 and 455 of CHIKV HVD. iii) FHL1- and CD2AP-binding sites in CHIKV HVD partially overlap, but this does not preclude formation of a HVD/CD2AP/FHL1 triple complex. iv) HVD-FHL1 interaction is critically determined by LIM1 domain of FHL1. v) The F429F430 di-peptide in CHIKV HVD is a critical determinant of FHL1/HVD interaction. vi) Interaction of FHL1 with its binding site induces allosteric changes in the downstream, C-terminal aa sequence of CHIKV HVD. vii) Since FHL1 is abundant in muscle cells, alterations of FHL1-HVD by mutations in the CHIKV HVD may improve the safety of live attenuated vaccine candidates.

## ACKNOWLEDGMENTS

This study was supported by Public Health Service grants R01AI133159 and R01AI118867 to E.I.F. and R21AI146969 to I.F and Swedish Foundation for Strategic Research grant ITM17-0218 to P.A.

## Notes

### Competing Interest Statement

The authors have declared no competing interest.

